# Pathogenic *DDX3X* mutations impair RNA metabolism and neurogenesis during fetal cortical development

**DOI:** 10.1101/317974

**Authors:** Ashley L. Lennox, Ruiji Jiang, Lindsey Suit, Brieana Fregeau, Charles J. Sheehan, Kimberly A. Aldinger, Ching Moey, Iryna Lobach, Ghayda Mirzaa, Alexandra Afenjar, Dusica Babovic-Vuksanovic, Stéphane Bézieau, Patrick R. Blackburn, Jens Bunt, Lydie Burglen, Perrine Charles, Brian H.Y. Chung, Benjamin Cogné, Suzanne DeBrosse, Nataliya Di Donato, Laurence Faivre, Delphine Héron, A. Micheil Innes, Bertrand Isidor, Bethany L. Johnson-Kerner, Boris Keren, Amy Kimball, Eric W. Klee, Paul Kuentz, Sébastien Küry, Dominique Martin-Coignard, Cyril Mignot, Noriko Miyake, Caroline Nava, Mathilde Nizon, Diana Rodriguez, Lot Snijders Blok, Christel Thauvin-Robinet, Julien Thevenon, Marie Vincent, Alban Ziegler, William Dobyns, Linda J. Richards, A. James Barkovich, Stephen N. Floor, Debra L. Silver, Elliott H. Sherr

**Affiliations:** Department of Molecular Genetics and Microbiology, Duke University Medical Center, Durham, NC 27710, USA; Department of Neurology, University of California, San Francisco, San Francisco, CA 94158, USA; Center for Integrative Brain Research, Seattle Children’s Research Institute, Seattle, WA 98101, USA; The University of Queensland, Queensland Brain Institute, Brisbane, 4072, Australia; Department of Epidemiology and Biostatistics, University of California San Francisco, San Francisco, CA, 94158, USA; APHP, Centre de référence des malformations et maladies congénitales du cervelet Département de génétique et embryologie médicale, Sorbonne Université, GRC n°19, pathologies Congénitales du Cervelet-LeucoDystrophies, AP-HP, Hôpital Armand Trousseau, F-75012 Paris, France; Department of Laboratory Medicine and Pathology, Mayo Clinic, Rochester, MN, USA; Department of Clinical Genomics, Mayo Clinic, Rochester, MN, USA; Department of Pediatric and Adolescent Medicine, Mayo Clinic, Rochester, MN, USA; CHU Nantes, Service de Génétique Médicale, 9 quai Moncousu, 44093, Nantes, CEDEX 1, France; l’institut du thorax, INSERM, CNRS, UNIV Nantes, 44007 Nantes, France; Center for Individualized Medicine, Mayo Clinic, Rochester, MN, USA; Department of Health Sciences Research, Mayo Clinic, Rochester, MN, USA; Centre de référence des Malformations et maladies congénitales du cervelet and Département de Génétique et embryologie médicales, AP-HP, GHUEP, Hôpital Trousseau 75012 Paris, France; Sorbonne Université, GRC n°19, Pathologies Congénitales du Cervelet LeucoDystrophies, APHP, Hôpital Armand Trousseau, F-75012 Paris, France; APHP, Département de Génétique, Centre de Référence Déficiences Intellectuelles de Causes Rares, Groupe Hospitalier Pitié Salpêtrière et GHUEP Hôpital Trousseau; Sorbonne Université, GRC “Déficience Intellectuelle et Autisme” Paris, France; Department of Paediatrics and Adolescent Medicine, Li Ka Shing Faculty of Medicine, The University of Hong Kong, Hong Kong, China; Center for Human Genetics, University Hospitals, Case Medical Center, Cleveland, OH 44195, USA; Institute for Clinical Genetics, TU Dresden, Dresden, Germany; Centre de référence Anomalies du Développement et Syndromes Malformatifs, INSERM UMR 1231 GAD, CHU de Dijon et Université de Bourgogne, Dijon, France; Department of Human Genetics, Yokohama City University Graduate School of Medicine, Yokohama 236-0004, Japan; Department of Medical Genetics, University of Calgary, Calgary, Alberta, Canada; APHP, Département de Génétique, Groupe Hospitalier Pitié Salpêtrière, Paris, France; Harvey Institute of Human Genetics, Greater Baltimore Medical Center, Baltimore, MD, USA; UMR-Inserm 1231 GAD, Génétique des Anomalies du développement, Université de Bourgogne Franche-Comté, Dijon, France; Service de Génétique, Centre hospitalier du Mans, Le Mans, France; APHP, GHUEP, Hôpital Armand Trousseau, Service de Neurologie Pédiatrique, Paris, France; Department of Human Genetics, Radboud University Medical Center, 6500 HB Nijmegen, the Netherlands; Centre de référence Déficience Intellectuelle, INSERM UMR 1231 GAD, CHU de Dijon et Université de Bourgogne, Dijon, France; Service de Génétique, CHU d’Angers, Angers, France; Departments of Pediatrics and Neurology, University of Washington, Seattle, WA 98101, USA; The University of Queensland, School of Biomedical Sciences, Brisbane, 4072, Australia; Department of Radiology and Biomedical Imaging, University of California, San Francisco, San Francisco, CA, USA; Department of Cell and Tissue Biology, UCSF, San Francisco, CA 94158 USA; Helen Diller Family Comprehensive Cancer Center, San Francisco, CA, 94158 USA; Department of Cell Biology, Duke University Medical Center, Durham, NC 27710 USA; Department of Neurobiology, Duke University Medical Center, Durham, NC 27710 USA; Duke Institute for Brain Sciences, Duke University, Durham, NC 27710 USA; Institute of Human Genetics and Weill Institute for Neurosciences, UCSF, SF CA 94158 USA

**Keywords:** polymicrogyria, corpus callosum agenesis, DDX3X, cortical development, radial glial progenitor, migration, stress granule, helicase

## Abstract

*De novo* germline mutations in the RNA helicase *DDX3X* account for 1-3% of unexplained intellectual disability (ID) cases in females, and are associated with autism, brain malformations, and epilepsy. Yet, the developmental and molecular mechanisms by which *DDX3X* mutations impair brain function are unknown. Here we use human and mouse genetics, and cell biological and biochemical approaches to elucidate mechanisms by which pathogenic *DDX3X* variants disrupt brain development. We report the largest clinical cohort to date with *DDX3X* mutations (n=78), demonstrating a striking correlation between recurrent dominant missense mutations, polymicrogyria, and the most severe clinical outcomes. We show that *Ddx3x* controls cortical development by regulating neuronal generation and migration. Severe *DDX3X* missense mutations profoundly disrupt RNA helicase activity and induce ectopic RNA-protein granules and aberrant translation in neural progenitors and neurons. Together, our study demonstrates novel mechanisms underlying *DDX3X* syndrome, and highlights roles for RNA-protein aggregates in the pathogenesis of neurodevelopmental disease.

## Introduction

From previous reports, it is estimated that between 1-3% of females with unexplained intellectual disability (ID) may have *de novo* nonsense, frameshift, splice site or missense mutations in *DDX3X* (Snijders Blok et al., 2015) and (https://doi.org/10.1101/283598). In these publications, approximately 40 individuals have been reported who present with diverse neurologic phenotypes, including microcephaly, corpus callosum hypoplasia, ventricular enlargement, and epilepsy, which together are identified as the *DDX3X* syndrome*. De novo* mutations in *DDX3X* have also been identified in individuals with autism spectrum disorder (ASD) (Iossifov et al., 2014; RK et al., 2017) and Toriello-Carey syndrome, a causally heterogeneous disorder characterized by severe ID, microcephaly, agenesis of the corpus callosum, post-natal growth defects, and craniofacial abnormalities (Dikow et al., 2017; Toriello et al., 2016). In addition to its role in neurodevelopmental disorders, *DDX3X* is recurrently mutated in several cancers, including medulloblastoma and lymphoma (Jiang et al., 2015; Jones et al., 2012; Pugh et al., 2012; Robinson et al., 2012) and in some cases identical residues are mutated in both ID and cancer. Early studies linked missense and nonsense mutations mechanistically by suggesting that *DDX3X* missense mutations may function primarily in a haploinsufficient manner through WNT signaling (Snijders Blok et al., 2015). However, to understand the likely pathophysiology, these phenotype/genotype correlations require detailed investigation with a large and fully phenotyped cohort of patients.

*DDX3X* encodes an RNA-binding protein of the DEAD-box family (Sharma and Jankowsky, 2014). While broadly implicated in mRNA metabolism, DDX3X is best characterized as a translational regulator (Lai et al., 2008; Shih et al., 2008). DDX3X is a component of RNA-protein granules, including neuronal transport granules (Elvira et al., 2006; Kanai et al., 2004) and cytoplasmic stress granules, which are RNA-protein foci induced by stress (Markmiller et al., 2018). DDX3X is not only required for assembly of stress granules (Shih et al., 2012), but overexpression of the wild-type (WT) protein or expression of cancer-associated mutations is sufficient to nucleate stress granules (Lai et al., 2008; Valentin-Vega et al., 2016). However, it is unknown whether aberrant RNA-protein granules induced by DDX3X contribute to neurodevelopmental pathologies of *DDX3X* syndrome.

In animal models, *Ddx3x* is essential for cell viability and division. *Ddx3x* depletion from mouse zygotes impairs blastocyst divisions and induces apoptosis, while depletion from immortalized cells disrupts chromosome segregation (Li et al., 2014; Pek and Kai, 2011). Likewise, in *Drosophila* germline stem cells, the *DDX3X* ortholog, *Belle*, is required for mitotic progression and survival (Kotov et al., 2016). Germline *Ddx3x* mouse mutant embryos exhibit apoptosis, DNA damage and cell cycle arrest, resulting in lethality at embryonic day (E) 9.5, along with aberrant neural tube closure (Chen et al., 2016). Notably, these phenotypes are present only in *Ddx3x* hemizygous male mice, whereas *Ddx3x* heterozygous female mice are viable. *DDX3X* is located in a 735 kb region in Xp11.4 that contains a small cluster of four genes (*CXorf38, MED14, USP9X, DDX3X*) that escape X inactivation. Several studies have shown that escape for DDX3X is developmentally regulated, tissue specific, and individually highly variable involving 30-60% of cells (Chen et al., 2016) and (https://www.biorxiv.org/content/early/2018/04/11/298984).

*DDX3X* brain malformations overwhelmingly affect the cerebral cortex, and thus are likely rooted in defective development of this structure. Embryonic cortical development relies upon precise generation and migration of both excitatory and inhibitory neurons. The main glutamatergic neural precursors are radial glial cells (RGCs), which divide in the ventricular zone (VZ) to produce either neurons or basal progenitors. In mice, the primary basal progenitors are intermediate progenitors (IPs) which generate neurons in the sub-ventricular zone (SVZ). Neurogenesis occurs between E11.5 and E18.5 in mice. Excitatory neurons are produced in an inside-out fashion, and following their genesis, migrate into the cortical plate (CP) using the RGC basal process as a scaffold (Taverna et al., 2014). Disorganization of the RGC basal process and/or defective neuronal migration are hypothesized to cause polymicrogyria (PMG) (Jamuar and Walsh, 2015). Likewise, aberrant neuronal generation underlies microcephaly. Although DDX3X function in cortical development has yet to be examined, it is essential for neurite outgrowth of cultured rat cortical and hippocampal neurons (Chen et al., 2016). Together with the cell cycle studies highlighted above, this suggests that defects in neural progenitor differentiation or neuronal migration, as well as disrupted formation of the radial glial scaffold, could underlie brain malformations associated with *DDX3X* syndrome.

While *DDX3X* mutations have been clearly linked to profound clinical deficits, the developmental and molecular mechanisms, as well as critical genotype-phenotype correlations, remain almost entirely unknown (Figure 1A). Here we address these essential issues by demonstrating that *DDX3X* mutations perturb RNA metabolism, resulting in aberrant development of the cerebral cortex in mice and humans. We identify 78 individuals with missense, frameshift, nonsense and splice-site *DDX3X* mutations, including nine recurrently mutated conserved residues. 76 of these mutations are in females and not found in either parent (hence *de novo*) and two male carriers harbor inherited hemizygous *DDX3X* mutations. Ten *DDX3X* mutations result in more severe clinical deficits, including PMG and disrupted corpus callosum anatomy, than seen in patients harboring loss-of-function (LoF) mutations. We then use *in vivo* studies in mice to demonstrate that *Ddx3x* LoF impairs neural progenitor function, resulting in reduced neuronal generation and impaired neuronal migration in the developing cerebral cortex. Finally, we show that *DDX3X* missense mutations cause formation of aberrant RNA-protein granules and altered translation patterns in neural progenitors and neurons, likely due to reduced helicase activity and impaired RNA release. Our data demonstrate a consistent correlation between severity of biochemical and cell biological assays and clinical outcomes. This work sheds light on novel and fundamental developmental and cellular mechanisms that underlie the etiology of *DDX3X* syndrome.

**Figure 1.**
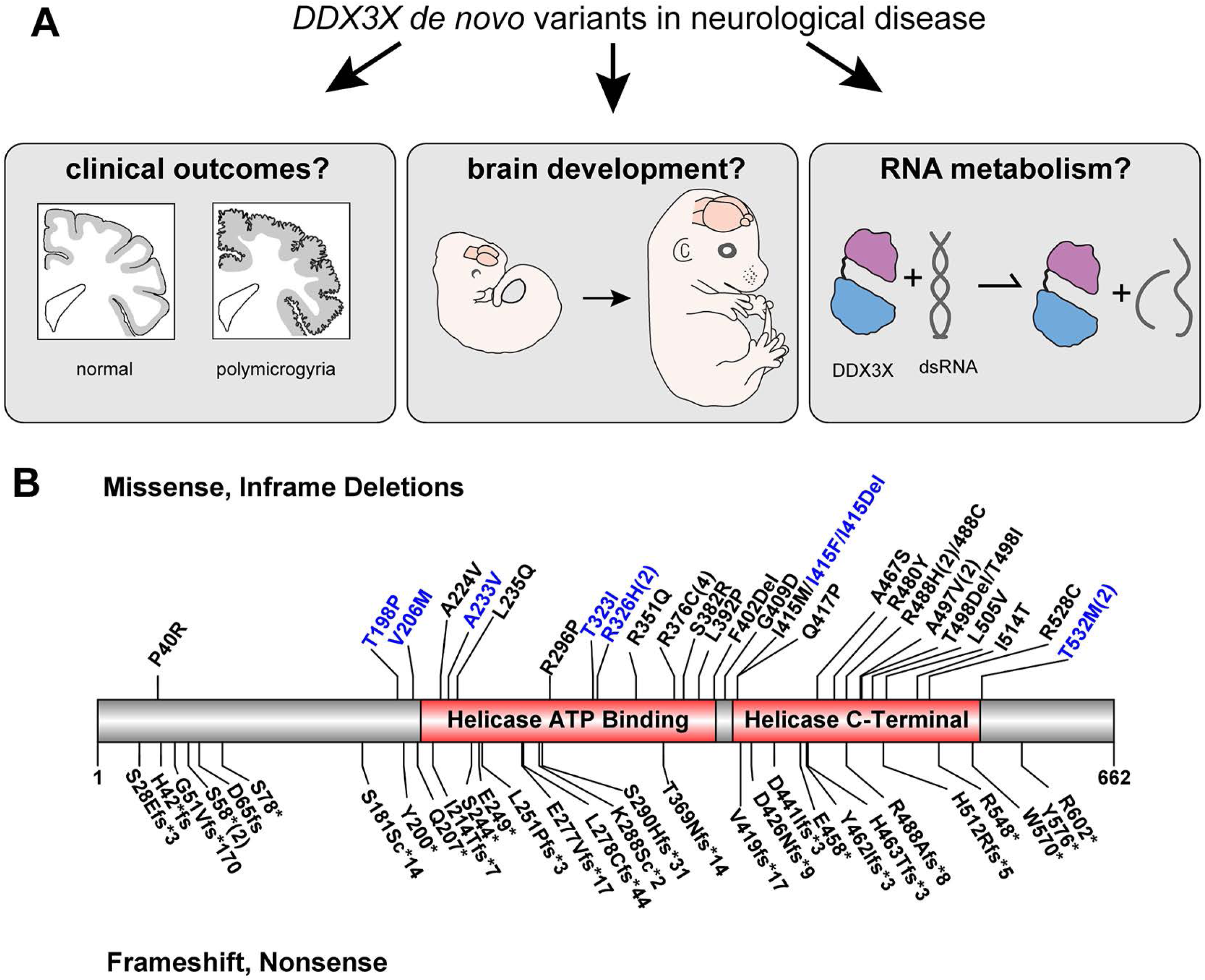
DDX3X Function and Predicted Amino Acid Changes in DDX3X in our Cohort. **A**, Overview of 3 questions assessed in this study to understand the role of *DDX3X de novo* mutations in disease. **B**, Missense and in frame deletions are annotated on top while frameshift and nonsense mutations which likely cause early termination are annotated on the bottom. Mutations found in patients with polymicrogyria are displayed in blue, seven of eight of which are missense mutations, and one is an in-frame deletion (I415Del). Recurrent mutations are indicated by numbers in the parentheses. Overall, mutations are enriched in the helicase domains at a rate higher than random chance (p<0.0001). Not shown are splice site mutations, found at G51, G89, K288 and A499 as the change to the protein sequence could not be reliably predicted.

## Results

### Identification of 78 individuals with *DDX3X* mutations and the associated clinical spectrum

To comprehensively investigate the spectrum of clinical phenotypes associated with *DDX3X* mutations, we identified *de novo* mutations in the *DDX3X* gene in 76 females and two male carriers (Kellaris et al., 2018) with maternally inherited hemizygous *DDX3X* mutations (Figure 1A and Table S1; 11 of these individuals were previously reported (Snijders Blok et al., 2015)). This cohort was recruited through collaboration with clinical geneticists and neurologists, clinical genetic laboratories, and through the DDX3X family support foundation (ddx3x.org).

To characterize the clinical features of these patients we obtained MRI scans and complete medical records on 64 individuals. 44 families were able to complete the Vineland Adaptive Behavior Scales that assessed their child’s development in a standardized manner. Two screening tools, the Social Responsiveness Scale-II (42 participants) and the Social Communication Questionnaire (36 participants) were used to assess risk for an ASD diagnosis and risk for social impairment (Figure S1). Within this cohort, 78/78 (100%) of all patients reported some neurologic finding, with the most common being ID followed by muscle tone abnormalities (62/72, 86%) (Table 1). Given that patients are clinically ascertained in this manner the penetrance is not unexpected, but the severity of neurologic and cognitive findings we observed does not automatically follow from this same ascertainment bias. We compared patient scores on the aforementioned standardized tests (Vineland, SRS-II, SCQ) to neurotypical controls. Overall, patients had a significant deviation in the mean score in all 3 exams (Vineland: 57.6 p<0.00001; SRSII 72.0, SCQ 16.7) when compared to neurotypical populations (100, <59, <15). Variation between scores on different tests was observed in the same patient, and a degree of variation equal to the variation observed in a control cohort was seen in the *DDX3X* mutation carriers (Figure S1, Table S1). The scores on the SRS-II and the SCQ suggest that 60% of the cohort score were above the “at risk” threshold for ASD and should be evaluated by a trained clinician to test whether the child’s clinical challenges meet criteria for ASD. Seventeen individuals (24%) in the cohort have seizures (ranging from Doose syndrome and infantile spasms to focal partial seizures or generalized absence spells), while 26 participants (41%) have microcephaly (defined as less than or equal to the 3^rd^ percentile). Other less common issues include cortical visual impairment, optic coloboma, strabismus, and both sensorineural and conductive hearing loss (Table 1). In addition to CNS related conditions, cardiac malformations are also present, including ventricular septal defects and atrial septal defects requiring repair. This is consistent with a prior case report suggesting that *DDX3X* mutation is associated with Toriello Carey syndrome (Dikow et al., 2017). Scoliosis was present in 10 participants (14%), which is likely linked to the significant hypotonia and weakness. Two patients were reported to have neuroblastoma, which were both detected incidentally. While previous literature on patients with germline *DDX3X* mutations focused primarily on the finding of intellectual and developmental disability (Snijders Blok et al., 2015), these data suggest that the full clinical spectrum found in these patients includes involvement outside the CNS, with significant implications for medical management.

**Table 1.**
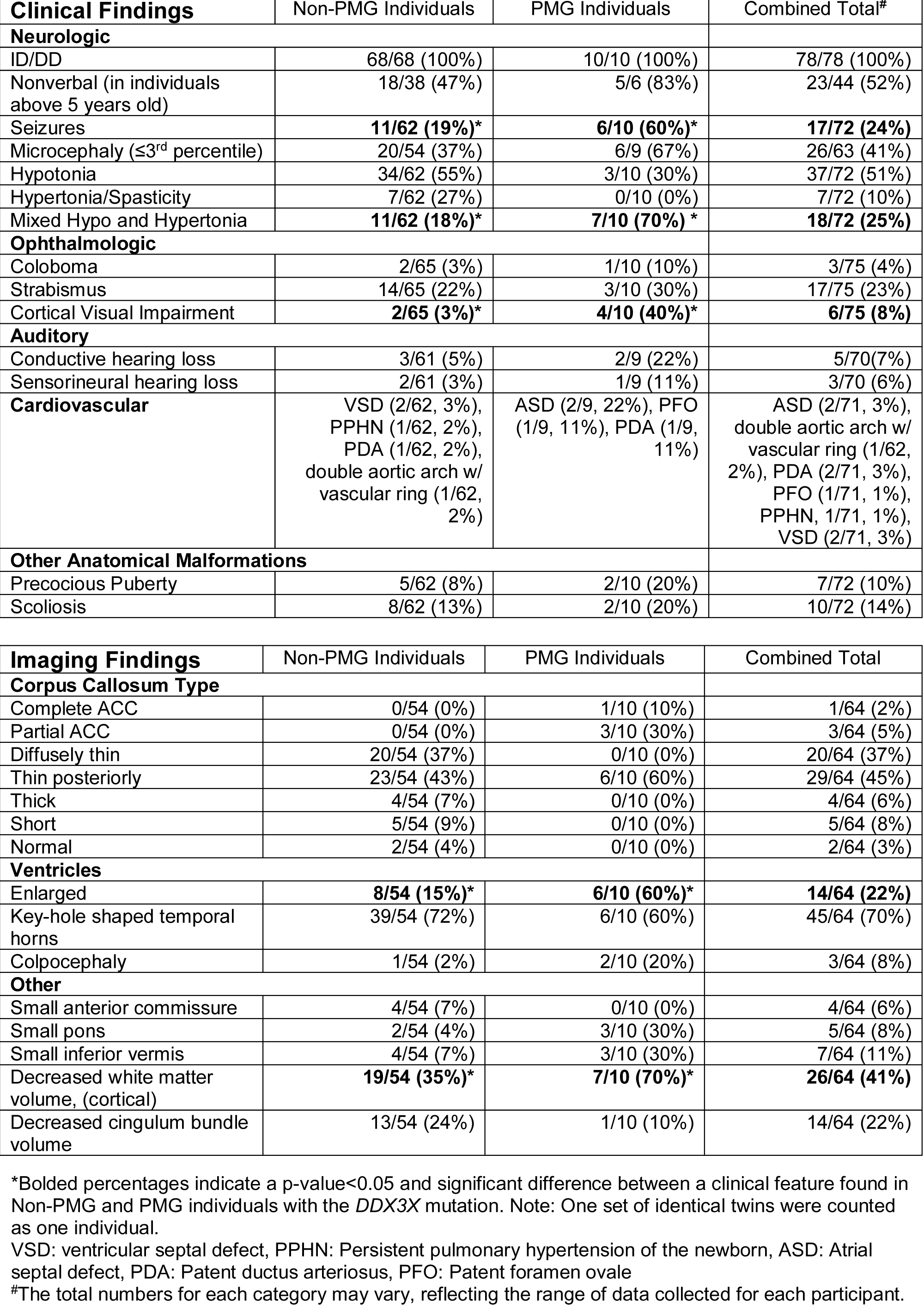
Clinical and Imaging Findings in Individuals with Mutations in *DDX3X*.

| **Clinical Findings** | Non-PMG Individuals | PMG Individuals | Combined Total<sup>#</sup> |
| --- | --- | --- | --- |
| **Neurologic** |  |  |  |
| ID/DD | 68/68 (100%) | 10/10 (100%) | 78/78 (100%) |
| Nonverbal (in individuals above 5 years old) | 18/38 (47%) | 5/6 (83%) | 23/44 (52%) |
| Seizures | **11/62 (19%)\*** | **6/10 (60%)\*** | **17/72 (24%)** |
| Microcephaly ($\leq 3^{\text{rd}}$ percentile) | 20/54 (37%) | 6/9 (67%) | 26/63 (41%) |
| Hypotonia | 34/62 (55%) | 3/10 (30%) | 37/72 (51%) |
| Hypertonia/Spasticity | 7/62 (27%) | 0/10 (0%) | 7/72 (10%) |
| Mixed Hypo and Hypertonia | **11/62 (18%)\*** | **7/10 (70%)\*** | **18/72 (25%)** |
| **Ophthalmologic** |  |  |  |
| Coloboma | 2/65 (3%) | 1/10 (10%) | 3/75 (4%) |
| Strabismus | 14/65 (22%) | 3/10 (30%) | 17/75 (23%) |
| Cortical Visual Impairment | **2/65 (3%)\*** | **4/10 (40%)\*** | **6/75 (8%)** |
| **Auditory** |  |  |  |
| Conductive hearing loss | 3/61 (5%) | 2/9 (22%) | 5/70 (7%) |
| Sensorineural hearing loss | 2/61 (3%) | 1/9 (11%) | 3/70 (6%) |
| **Cardiovascular** | VSD (2/62, 3%),<br>PPHN (1/62, 2%),<br>PDA (1/62, 2%),<br>double aortic arch w/<br>vascular ring (1/62,<br>2%) | ASD (2/9, 22%), PFO<br>(1/9, 11%), PDA (1/9,<br>11%) | ASD (2/71, 3%),<br>double aortic arch w/<br>vascular ring (1/62,<br>2%), PDA (2/71, 3%),<br>PFO (1/71, 1%),<br>PPHN, 1/71, 1%),<br>VSD (2/71, 3%) |
| **Other Anatomical Malformations** |  |  |  |
| Precocious Puberty | 5/62 (8%) | 2/10 (20%) | 7/72 (10%) |
| Scoliosis | 8/62 (13%) | 2/10 (20%) | 10/72 (14%) |

| **Imaging Findings** | Non-PMG Individuals | PMG Individuals | Combined Total |
| --- | --- | --- | --- |
| **Corpus Callosum Type** |  |  |  |
| Complete ACC | 0/54 (0%) | 1/10 (10%) | 1/64 (2%) |
| Partial ACC | 0/54 (0%) | 3/10 (30%) | 3/64 (5%) |
| Diffusely thin | 20/54 (37%) | 0/10 (0%) | 20/64 (37%) |
| Thin posteriorly | 23/54 (43%) | 6/10 (60%) | 29/64 (45%) |
| Thick | 4/54 (7%) | 0/10 (0%) | 4/64 (6%) |
| Short | 5/54 (9%) | 0/10 (0%) | 5/64 (8%) |
| Normal | 2/54 (4%) | 0/10 (0%) | 2/64 (3%) |
| **Ventricles** |  |  |  |
| Enlarged | **8/54 (15%)\*** | **6/10 (60%)\*** | **14/64 (22%)** |
| Key-hole shaped temporal horns | 39/54 (72%) | 6/10 (60%) | 45/64 (70%) |
| Colpocephaly | 1/54 (2%) | 2/10 (20%) | 3/64 (8%) |
| **Other** |  |  |  |
| Small anterior commissure | 4/54 (7%) | 0/10 (0%) | 4/64 (6%) |
| Small pons | 2/54 (4%) | 3/10 (30%) | 5/64 (8%) |
| Small inferior vermis | 4/54 (7%) | 3/10 (30%) | 7/64 (11%) |
| Decreased white matter volume, (cortical) | **19/54 (35%)\*** | **7/10 (70%)\*** | **26/64 (41%)** |
| Decreased cingulum bundle volume | 13/54 (24%) | 1/10 (10%) | 14/64 (22%) |

\*Bolded percentages indicate a p-value<0.05 and significant difference between a clinical feature found in Non-PMG and PMG individuals with the *DDX3X* mutation. Note: One set of identical twins were counted as one individual.
VSD: ventricular septal defect, PPHN: Persistent pulmonary hypertension of the newborn, ASD: Atrial septal defect, PDA: Patent ductus arteriosus, PFO: Patent foramen ovale
#The total numbers for each category may vary, reflecting the range of data collected for each participant.

### *DDX3X* mutations impair human brain development

To identify the brain anatomic disruptions notable in our cohort of *DDX3X* mutation carriers, we obtained and systematically reviewed high quality brain MRI scans from 64 patients (Figure 2 and Table 1). Of these patients, 62/64 (97%) had a corpus callosum malformation, ranging from complete agenesis (1/64, 2%) to a milder impairment with only a thin posterior body and splenium (44/64, 69%). A small but significant percentage of patients (10/64, 16%) also had polymicrogyria (PMG), a condition characterized by abnormally dense and small gyri in the cerebral cortex (Figure 2D-F). Other noted features were globally diminished white matter volume (26/64, 41%), and lateral ventricles with a characteristic enlargement specifically in the temporal horns of the lateral ventricles, referred to as “key-hole-shaped” temporal horns (45/64, 70%). This finding is also evident in other cases of agenesis of the corpus callosum where it has been shown to correlate closely with diminished size of the ventral aspect of the cingulum bundles (Nakata et al., 2009) (Figure 2A1-C1). Other findings such as a diminished anterior commissure, small pons, and small inferior cerebellar vermis were present in a small subset of patients (Table 1). Thus, in the vast majority of *DDX3X* patients, mutations result in a range of brain anatomy changes, suggesting these findings are fundamentally and consistently linked to changes in brain development, which we address below.

**Figure 2.**
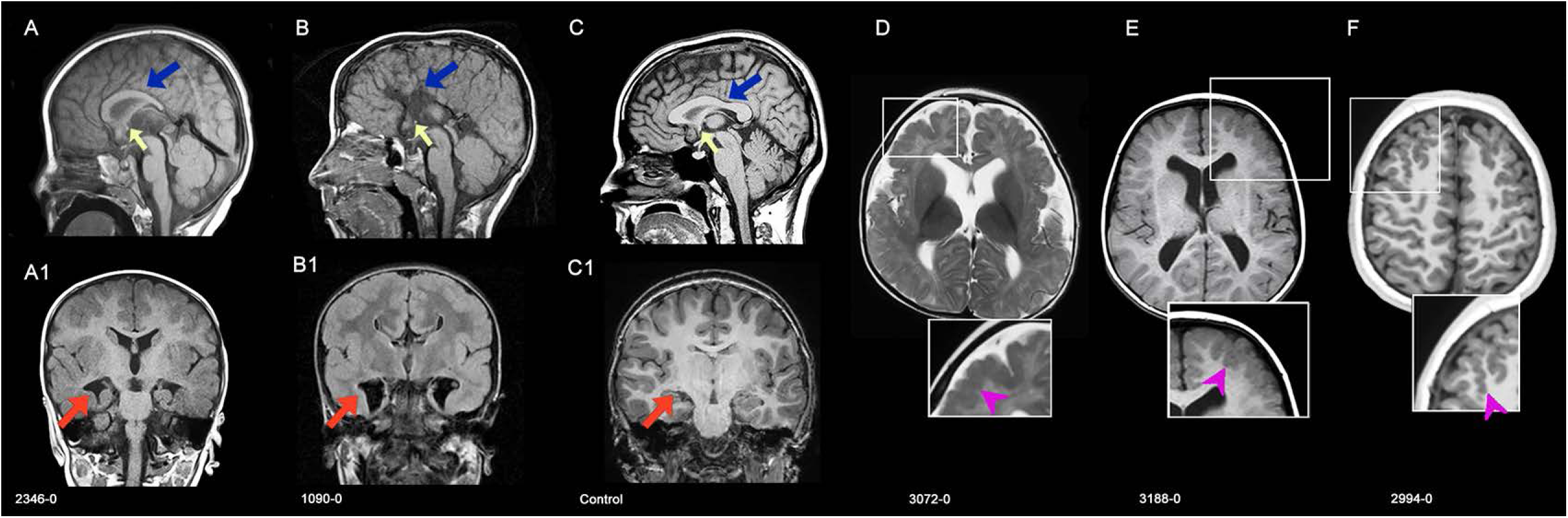
Common Brain Imaging Findings in DDX3X Cohort. **A,A1** Patient 2346–0 at 12 months. **A,** Sagittal image shows corpus callosum is rounded and hypogenetic with a short, thinned posterior body (blue arrow) and absent splenium; inferior genu and rostrum are absent. A small anterior commissure (yellow arrow) is present. **A1,** Coronal image shows hippocampi are abnormally rotated, including keyhole-shaped temporal horns with a narrow inferior extension (red arrow). Third ventricle is slightly enlarged. White matter volume was mildly decreased. Note abnormally thick cortex with too few, too shallow sulci; this has features of a mild cobblestone cortex. **B, B1,** Patient 1090-0 at 9 years of age. **A,** Sagittal image shows complete agenesis of the corpus callosum and hippocampal commissure (blue arrow), a small anterior commissure (yellow arrow), and colpocephaly with enlarged keyhole-shaped temporal horns (red arrow) that extend inferomedially most likely due to markedly hypoplastic ventral cingulum. **B1,** The cerebral cortex is too thick with irregular inner and outer surfaces, likely resulting from overmigration of neurons through gaps in the glia limitans (cobblestone cortex). White matter volume is diminished. **C, C1,** Normal MRIs for comparison. **C,** Mid-sagittal T1-weighted image showing a normal sized corpus callosum (blue arrow) and anterior commissure (yellow arrow). **C1,** Coronal T1-weighted image shows normal cortical thickness and gyration, normal sized ventricle bodies and temporal horns (red arrow). **D,** Patient LR14-020 at 6 months of age had extensive, asymmetric frontal and perisylvian cobblestone cortical dysgenesis, right worse than left (pink arrow head) and enlarged ventricles. **E,** Patient LR07-255 at 7 years of age also had extensive frontal cobblestone cortical dysgenesis, left (pink arrow head) greater than right, and enlarged ventricles. **F,** Patient 2994-0 at 10 years of age had bilateral frontal cobblestone cortical dysgenesis (pink arrowhead).

### Location and type of mutation predicts imaging features and clinical outcomes

We sought to determine if the location and type of *DDX3X* mutation found in patients correlated with the severity of their clinical phenotype. We first focused on mutation type, for which there are 38 individuals with missense mutations or in-frame deletions, 32 individuals with nonsense or frameshift mutations (LoF), and eight patients with splice site mutations (Table S1). Patients with missense mutations were significantly more likely to have a severely impaired radiological or clinical phenotype, such as PMG (p=0.001) and cortical visual impairment (CVI) (p=0.007) as compared to the cohort overall. In parallel, not a single patient with a frameshift or nonsense LoF mutation presented with PMG (p=0.0004), and the prevalence of CVI was also lower in the patients with LoF mutations (nearly statistically significant (p=0.054) than in the overall group.

The high rate of recurrent mutations led us to ask whether patients with identical mutations present with similar clinical phenotypes. In the cohort we identified nine nucleotide positions that were repeatedly mutated; and the probability of each of these individual loci having recurrent mutations, given the number of overall mutations, was low and statistically significant for each grouping. This includes four patients each with mutations at R376 or A488 (p<1x10^-7^), three patients with mutations at I415 (p=1.13e-05), and two patients each with mutations at the remaining 6 recurrent loci (p=0.00083). Of the loci with recurrent mutations, while the numbers were too small to conduct statistical analyses comparing the severity of clinical deficits, qualitative analysis suggests that they result in very similar phenotypes. For example, two patients who each had the same *de novo* mutation at R326H exhibited a similarly severe imaging and clinical phenotype, notable for PMG, microcephaly, severe developmental delay, and optic nerve hypoplasia. Conversely, the four patients with the R376C mutation presented with no cortical malformations, a thin corpus callosum body and splenium, and a generally less severe clinical presentation with 3 out of 4 patients speaking in full sentences, having an average Vineland score of 64, and none having seizures. Non-neurologic phenotypes can also cluster, as 5/9 patients with cardiac malformations (VSD, ASD, PFO, PPHN, aortic arch irregularities) had mutations at just two recurrent loci (I415, R488).

Strikingly, of the 10 patients who had PMG, nine had missense mutations (one had an in-frame single amino acid deletion) with 6 mutations found at three recurrent loci (T532, I415, R326). This suggests that these recurrently mutated loci are more commonly associated with PMG (χ^2^=5.14; p<0.023). PMG patients presented with a more severe anatomic brain phenotype as 6 of 9 (67%) had microcephaly (<3^rd^ percentile), whereas only 20 of the 54 individuals (35%) without PMG had microcephaly. This suggests a potentially mechanistic linkage between microcephaly and PMG in this cohort. This association between missense mutations and PMG suggests that this subset of missense mutations function in a dominant manner, more severe than any of the patients with LoF mutations, none of whom have PMG. Furthermore, 3/9 patients with cardiac findings (VSD, ASD, aortic abnormalities) also had PMG, disproportionate to the number of PMG patients in the overall cohort. PMG patients were also more likely to have cortical visual impairment (40% vs 3% p<0.0001), seizures (60% vs 19% p=0.007), and partial or complete ACC (40% vs 0% p<0.0001). Overall, these data suggest that there is an association between certain clinical features and the type of mutation, with a subset of missense mutations being more likely to result in a severe clinical phenotype.

### *Ddx3x* is expressed in the developing mouse neocortex

The cortical and neurological findings we have described in *DDX3X* patients suggest this gene plays a central role in brain development, and, in particular, the cerebral cortex. In line with this hypothesis, *DDX3X* is expressed across human fetal brain regions from post conception weeks (PCW) 11-22 (Miller et al., 2014), and in cortical progenitors and neurons by single cell sequencing of human fetal brain samples (Nowakowski et al., 2017). Likewise, genomic studies of the developing mouse neocortex show *DDX3X* is highly expressed in both progenitors and neurons, with especially high expression in newborn callosal projection neurons (Ayoub et al., 2011; Molyneaux et al., 2009; Molyneaux et al., 2015). DDX3X is highly conserved, showing 98.6% identity at the amino acid level between mice and humans. Taken together, this suggests that mice are a suitable model for investigating DDX3X LoF requirements for cortical development (Figure 3A).

**Figure 3.**
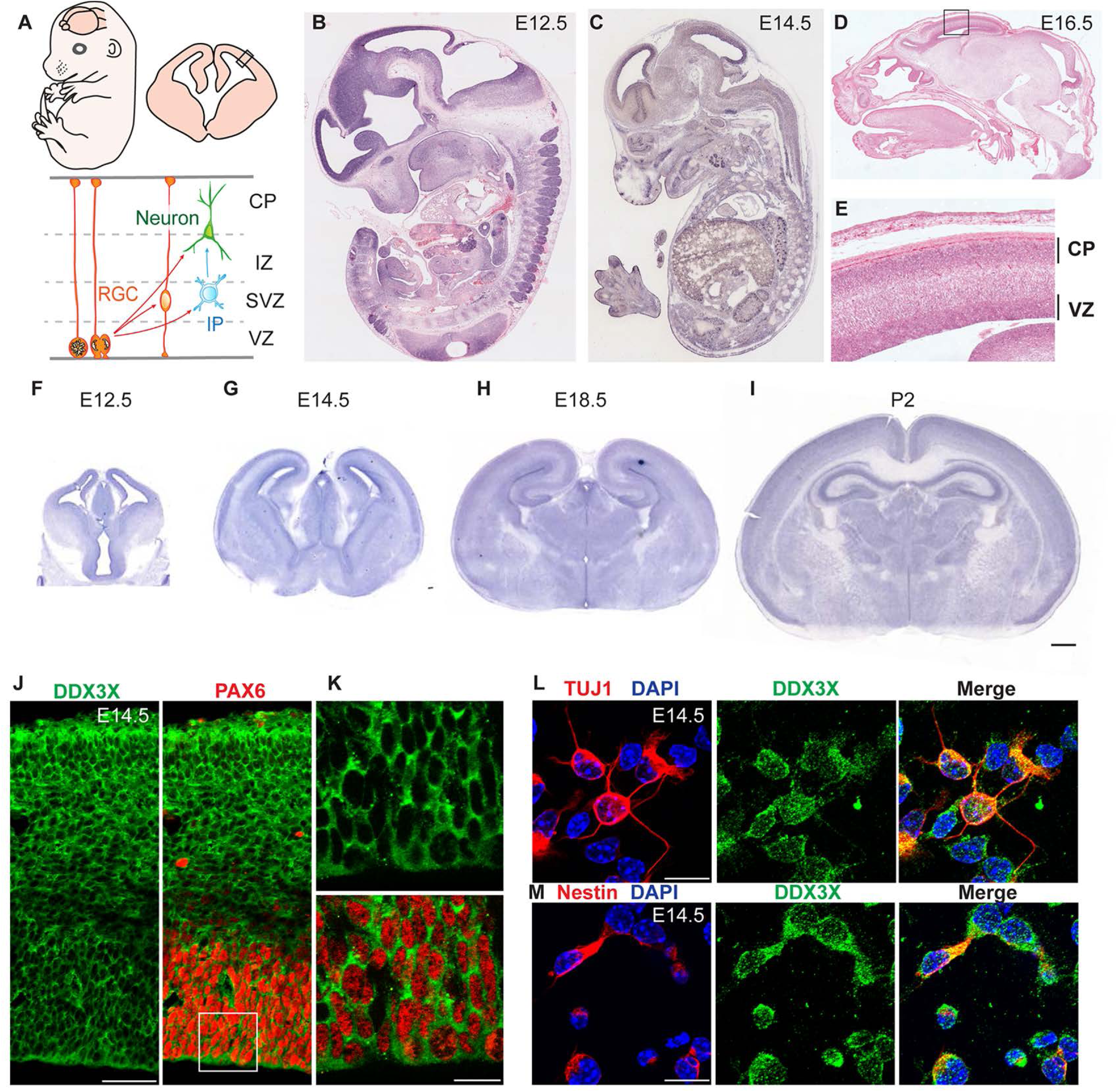
*Ddx3x* is expressed in progenitors and neurons in the embryonic mouse cortex. **A**, Top, diagram representing an embryonic mouse and a coronal section through the cortex with box highlighting region depicted in lower panel. Bottom, diagram representation of major cell types examined in this study including radial glial progenitors (RGC), intermediate progenitors (IP), and neurons. Zones of the cortex are demarcated with dotted lines demonstrating boundaries of ventricular zone (VZ), subventricular zone (SVZ), intermediate zone (IZ), and cortical plate (CP). **B-E,** *In situ* hybridization against *Ddx3x* in sagittal sections in E12.5 (**B**), E14.5 (**C**), and E16.5 (**D,E**) embryos. Box in **D** is magnified in **E**. **F-I**, *In situ* hybridization against *Ddx3x* mRNA in coronal sections in E12.5 (**F**), E14.5 (**G**), E18.5 (**H**), and P2 (**I**) brains. **J-K,** Immunofluorescence for DDX3X (green) in E14.5 cortical sections co-stained for the progenitor marker PAX6 (red). Boxed region in **J** is magnified in **K**. **L-M,** Immunofluorescence for DDX3X (green) in cultured primary cells isolated from E14.5 cortices co-stained for the neuron marker TUJ1 (**L**) or the progenitor marker Nestin (**M**) in red and DAPI in blue. Scale bars: 500 μm (**I**); 50 μm (**J**); 15 μm (**K, L, M**).

We first assessed *Ddx3x* spatial and temporal expression in mouse cortical development using *in situ* hybridization. *In situ* hybridization of E12.5 and E14.5 sagittal sections showed *Ddx3x* transcript is ubiquitous, but especially enriched in the developing mouse brain (Visel et al., 2004) (Figures 3B and C). Within the developing cerebral cortex, *Ddx3x* expression was initially enriched in progenitor populations of the VZ (Visel et al., 2004) (Figures 3B, C, F, G). By E15.5 onwards, *Ddx3x* mRNA expression was evident in both the CP and VZ/SVZ (Figures 3D, E, G, H). By postnatal day 2, *Ddx3x* mRNA was highly expressed throughout the pyramidal layers in the hippocampus (CA1-CA3 and subiculum) and minimally expressed in the corpus callosum (Figure 3I). This analysis suggests that *Ddx3x* is widely expressed in the brain, especially within cortical regions, and spanning mouse fetal cortical development.

We next evaluated DDX3X protein expression. Consistent with its RNA expression, DDX3X was expressed in all cortical layers in embryonic brains (Figures 3J and K). In primary cells, DDX3X was detected in both Nestin-positive progenitors and TUJ1-positive neurons (Figures 3L and M). Using markers for neurons, astrocytes, oligodendrocytes, microglia and ependymal cells, we detected DDX3X expression in all major cell types of P5 brains (Figure S2). Notably, neuronal cells had comparatively higher DDX3X expression compared to other cell types, consistent with embryonic expression in neural progenitors and neurons. In both cortical sections and cultured cells, DDX3X primarily localized to puncta within the cytoplasm (Figures 3J-M). This pattern is consistent with its known roles as an RNA helicase, and with previous descriptions of its subcellular localization in immortalized cells (Lai et al., 2008). Taken together with the genomic data illustrating DDX3X expression, this demonstrates that DDX3X is expressed throughout cortical development in both neurons and progenitors.

### *In vivo* disruption of *Ddx3x* alters corticogenesis

We next interrogated the requirement of *Ddx3x* in embryonic corticogenesis using CRISPR/Cas9 to create loss-of-function mutations in *Ddx3x*. We designed a short guide RNA (sgRNA) against Exon 1 of the *Ddx3x* gene approximately 20 nucleotides downstream of the start codon (Figure 4A). We reasoned that indels generated by double-strand break repair would cause frameshift mutations and premature stop codons early in the transcript, ultimately resulting in heterogeneous LoF mutations. We assessed the efficacy of *Ddx3x* Exon1 sgRNA in Neuro2A cells. Three-day expression of *Ddx3x* sgRNA + Cas9 led to a 50% and 60% reduction in *Ddx3x* mRNA and protein levels, respectively, relative to control (Cas9 without sgRNA) (Figures 4B-D). Notably, these experiments were carried out in an unsorted population of Neuro2A cells with approximately 60% transfection efficiency, suggesting the sgRNA effectively depletes *Ddx3x* levels on average by an estimated 90-100% in transfected cells.

**Figure 4.**
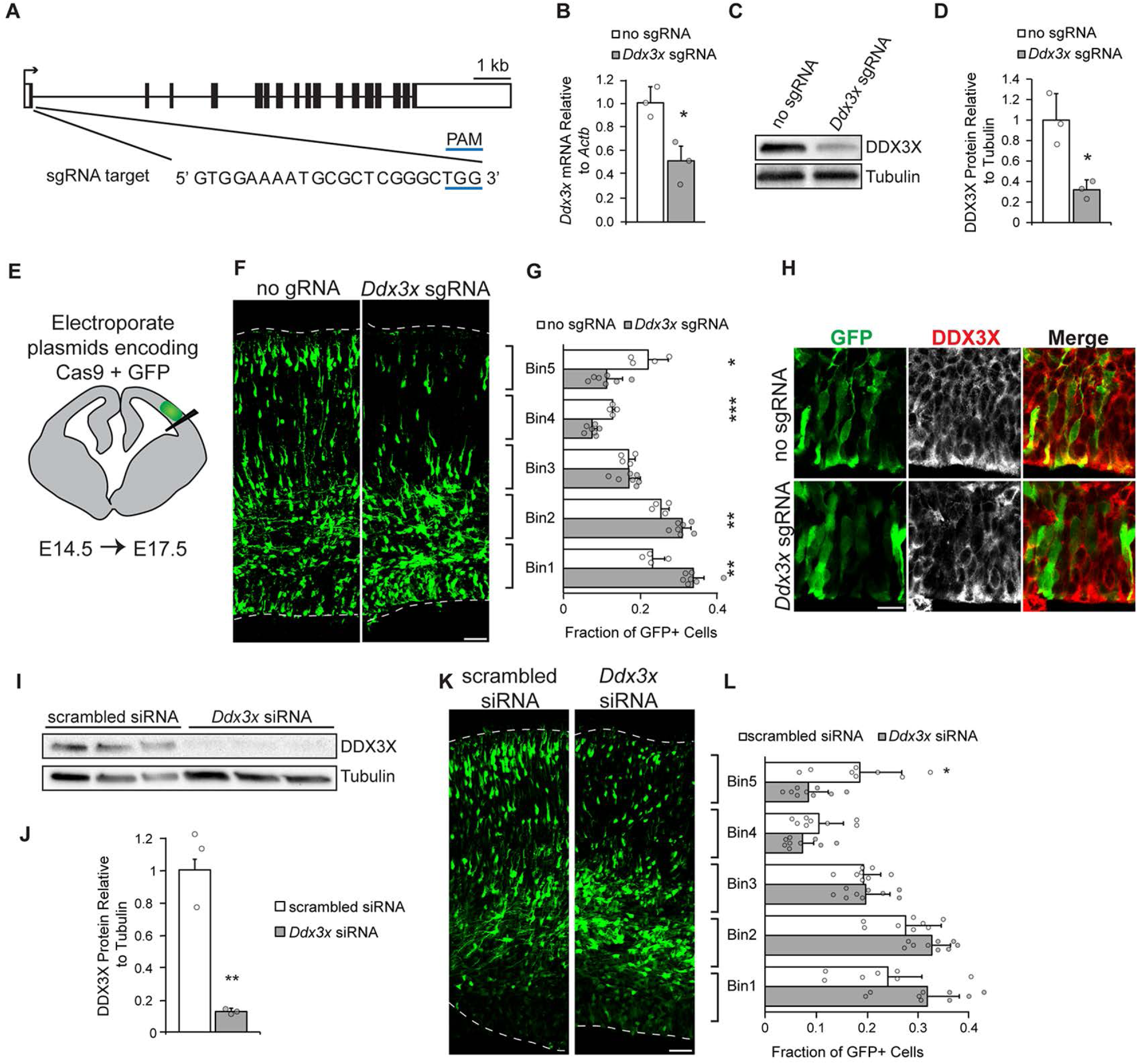
*In vivo* disruption of *Ddx3x* alters neurogenesis. **A,** Schematic of mouse *Ddx3x* gene structure with exons (boxes) and introns (thin lines) depicted. *Ddx3x* sgRNA targets Exon 1 and the sequence is shown. **B-D**, Validation of *Ddx3x* sgRNA efficacy in Neuro2A cells assessed by depletion of mRNA and protein levels 3 days after transfection. Approximately 60% of cells were transfected. **B,** qPCR of *Ddx3x* mRNA in exons 5-6 normalized to *Actb* (n=3 biological replicates/condition, Student’s t-test p=0.038). **C,** Representative western blot of Neuro2A cells treated with no sgRNA or *Ddx3x* sgRNA and probed for DDX3X and Tubulin. **D,** Quantification of western blots by densitometry, following normalization to Tubulin (n=3 biological replicates/condition, Student’s t-test, p=0.032). **E,** Schematic of coronal section depicting site of electroporation. **F,** Representative coronal sections of E17.5 brains *in utero* electroporated at E14.5 with pCAG-GFP and pX330-Cas9 no sgRNA or pX330-Cas9 *Ddx3x* sgRNA. Sections were stained with anti-GFP antibody (green). Dotted lines represent ventricular and pial surfaces, and brackets on the right refer to the bins. **G,** Quantitation of distribution of GFP-positive cells in five evenly spaced bins with Bin1 at the ventricle and Bin5 at the pia (n=4 embryos (no gRNA) or 7 embryos (*Ddx3x* gRNA), Student’s t-test p=0.0015(Bin1); 0.0053(Bin2); 0.9519(Bin3); 0.00001(Bin4); 0.0161(Bin5)). **H.** Immunofluorescence for DDX3X (white or red) and GFP (green) in sections from **E** focused in VZ. **I-J,** Validation of *Ddx3x* siRNA efficacy in Neuro2A cells assessed by western blot three days after transfection. Representative western blot for DDX3X and Tubulin (**I**). Quantification of western blots by densitometry, following normalization to Tubulin (n=3 biological replicates, Student’s t-test p=0.047). **K,** Representative coronal sections of E17.5 brains *in utero* electroporated at E14.5 with pCAG-GFP and scrambled or *Ddx3x* siRNAs stained with anti-GFP (green). **L,** Quantification of distribution of GFP-positive cells in five evenly spaced bins (n=8 embryos (scrambled), or 9 embryos (*Ddx3x*), Student’s t-test p=0.1251(Bin1); 0.0564 (Bin2); 0.8419(Bin3); 0.1518(Bin4); 0.0169(Bin5)). Scale bars: 50 μm (**F,K**);15 μm (**I**). Error bars = standard deviation.

We next used *in utero* electroporation to deliver a plasmid encoding *Ddx3x* sgRNA and Cas9 along with a GFP-encoding plasmid to radial glial progenitors of the developing embryonic brain (Figure 4E). This approach is ideal for reducing *Ddx3x* expression in both radial glial progenitors and their progeny (Saito, 2006). We targeted progenitors at E14.5 during mid-corticogenesis when *Ddx3x* expression is robust and progenitors are actively neurogenic. Brains were harvested three days later at E17.5, allowing time for progenitors to divide and differentiate and for newborn neurons to migrate to the CP. We then assessed the distribution of GFP-positive cells within five evenly spaced bins in 450 μm wide cortical columns (Figure 4F). GFP-positive cells in control brains were evenly distributed across cortical bins. In contrast, expression of *Ddx3x* sgRNA significantly altered their distribution resulting in a 1.4-fold increase in cells within the VZ/SVZ proliferative zones (note bins 1 and 2) and a 1.9-fold decrease in cells in the CP (note bins 4 and 5) relative to no sgRNA controls (Figure 4G). We consistently detected reduced DDX3X immunostaining in GFP-positive cells expressing sgRNA but not the control, with some cell-to-cell variation as predicted with CRISPR-mediated approaches (Figure 4H). To assess *Ddx3x* requirements with an independent methodology, we used siRNAs targeting *Ddx3x* 3’UTR. A three-day knockdown in Neuro2A cells showed the siRNAs effectively depleted 90% of DDX3X protein (Figures 4I-J). We then used *in utero* electroporation to introduce either scrambled or *Ddx3x* siRNAs into E14.5 brains and harvested brains at E17.5. *Ddx3x* siRNA-mediated depletion caused significantly more electroporated cells in proliferative zones and less in the CP compared to a scrambled control, phenocopying CRISPR experiments above (Figures 4K-L). Together, these findings demonstrate that *Ddx3x* is required for proper distribution of newborn cells in the neocortex, suggesting a requirement in neurogenesis.

### *In vivo* disruption of *Ddx3x* impacts progenitor differentiation and neuronal migration

The aberrant distribution of neurons in the CP in *Ddx3x* LoF could involve defective progenitor proliferation or differentiation, selective neuronal cell death in the CP, and/or defects in neuronal migration. To investigate these possibilities, we repeated CRISPR-mediated *Ddx3x* knockdown at E13.5 and analyzed brains at E15.5. A two-day paradigm was used to pinpoint cellular defects leading to aberrant distribution, and analysis at E15.5 enabled us to quantify progenitors, which are more abundant at this stage. *Ddx3x*-depletion significantly altered GFP-positive cell distribution at E15.5 (Figures 5A and B), leading to more cells in the proliferative zones and fewer in the CP. This is consistent with experiments in Figure 4 and extend the developmental requirement of *Ddx3x* to include E13.5-E17.5.

**Figure 5.**
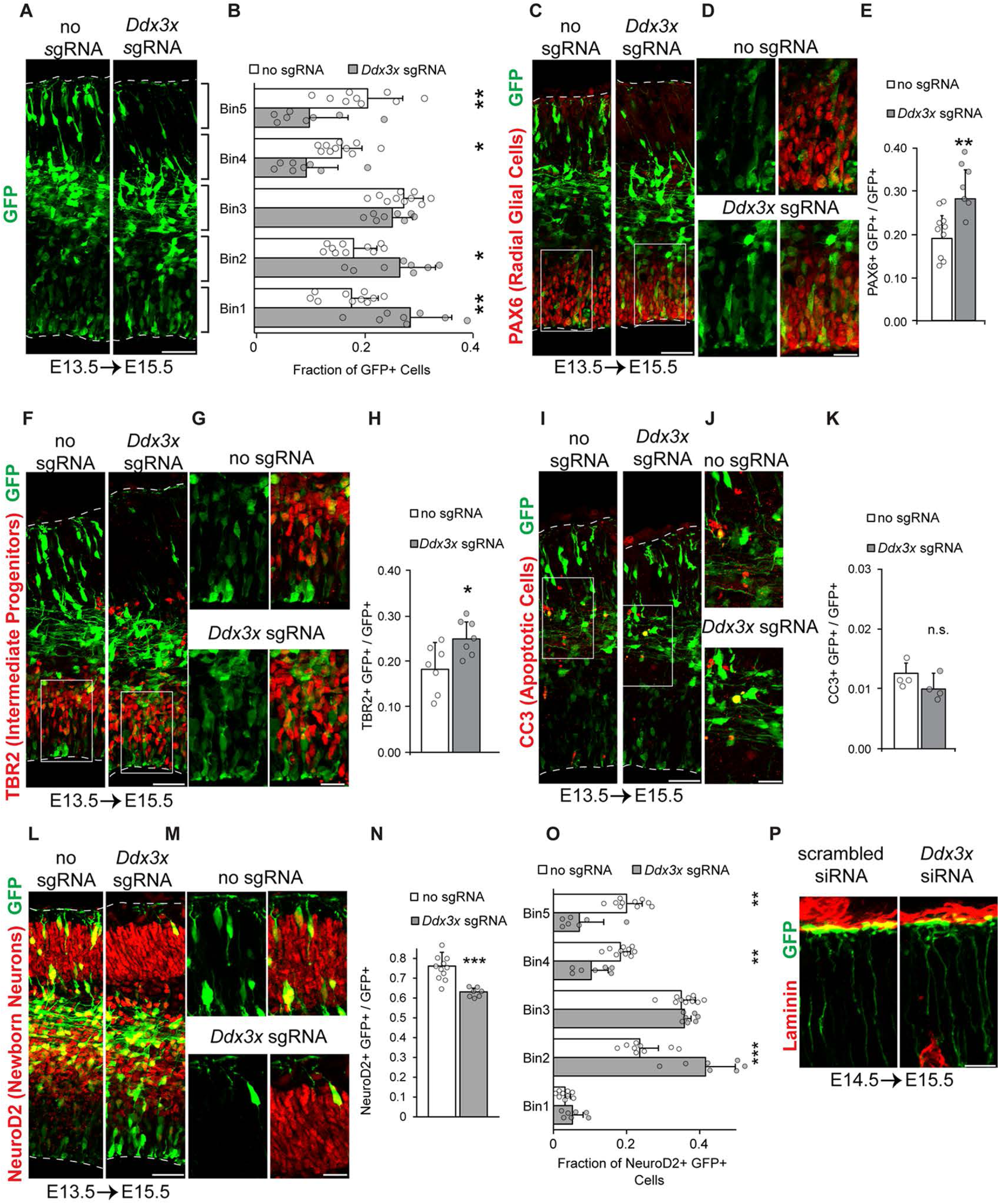
*Ddx3x* is required for neuron generation and neuron migration *in vivo*. **A,** Representative coronal sections of E15.5 brains *in utero* electroporated at E13.5 with pCAG-GFP and Cas9 no sgRNA or *Ddx3x* sgRNA stained with anti-GFP (green). **B,** Quantification of distribution of GFP-positive cells in **A** (n=9 embryos (no sgRNA) or 8 embryos (*Ddx3x* sgRNA), Student’s t-test p=0.005 (Bin1); 0.008 (Bin2); 0.249 (Bin3); 0.019 (Bin4); 0.005 (Bin5)). **C-D,** Sections of electroporated E13.5 – E15.5 brains stained with GFP (green) and PAX6 (red). Boxed regions in the VZ in **C** are shown at higher magnification in **D. E,** Quantification of percentage of GFP-positive cells expressing PAX6 (n=10 (no sgRNA) or 7 embryos (*Ddx3x* sgRNA), Student’s t-test p=0.007). **F-G,** Sections of electroporated E13.5 – E15.5 brains stained with GFP (green) and TBR2 (red). Boxed regions in the VZ in **F** are shown at higher magnification in **G. H,** Quantification of percentage of GFP-positive cells expressing TBR2 (n=7 embryos for each condition, Student’s t-test p=0.021). **I-J,** Sections of electroporated brains stained with GFP (green) and CC3 (red). Boxed regions in the VZ in **I** are shown at higher magnification in **J. K,** Quantification of percentage of GFP-positive cells expressing CC3 (n=4 embryos for each condition, Student’s t-test p=0.155). **L-M,** Sections of electroporated E13.5 – E15.5 brains stained with GFP (green) and NeuroD2 (red). Boxed regions in VZ in **L** are shown at higher magnification in **M. N**, Quantification of percentage of GFP-positive cells expressing NeuroD2 (n=10 (no sgRNA) or 7 embryos (*Ddx3x* sgRNA), Student’s t-test p=0.00016). **O,** Quantification of the distribution of GFP/NeuroD2 double-positive cells in **M** (Student’s t-test p=0.1295(Bin1); 0.0005 (Bin2); 0.5478 (Bin3); 0.0029 (Bin4); 0.0013 (Bin5)). **P**, Sections of E14.5 – E15.5 electroporations with siRNAs stained for GFP (green) and Laminin (red) to mark the basement membrane. Images are focused at the pial surface. Scale bars: 50 μm (**A, C, F, I, L**); 15 μm (**D, G, J, M, P**). Error bars = standard deviation.

We then quantified the impact of *Ddx3x* LoF upon progenitors and neurons. In GFP-positive cells, we measured expression of the transcription factors PAX6 and TBR2 to mark RGCs and IPs, respectively. Relative to the control, *Ddx3x* LoF led to a significant 1.5-fold increase in PAX6+ radial glial progenitors (Figures 5C-E, p=0.007) and a 1.4-fold increase in TBR2+ intermediate progenitors (Figures 5F-H, p=0.021). To assess neuron number, we quantified GFP-positive cells expressing the transcription factor NeuroD2. *Ddx3x*-deficiency in progenitors resulted in a significant 1.2-fold decrease in new neurons (Figures 5L-N, p=0.0002). Since reduced neurons are concomitant with increased progenitors, this suggests *Ddx3x* controls the normal balance of progenitors and differentiated neurons *in vivo.* To determine if this imbalance is due to selective neuronal cell death, we quantified cleaved-Caspase3 (CC3) (Figure 5I-K). However, we noted equivalent low levels of apoptosis in both mutant and control brains (p=0.1552). Together, these results indicate that *Ddx3x* is required for proper differentiation of progenitor cells.

After neurons are generated by progenitors in the VZ/SVZ, they migrate radially into the CP. Given that *Ddx3x* is also expressed in neurons, we monitored neuronal migration following *Ddx3x* depletion by quantifying the distribution of GFP and NeuroD2 double-positive cells within cortical bins (Figure 5O). Between E13.5-E15.5, 20% of control cells reached the upper CP, whereas only 7% of *Ddx3x-*depleted cells did (Figure 5O, Bin 5 p=0.00127). Similarly, between E14.5-E17.5, *Ddx3x* depleted neurons were also reduced in number with delayed migration to the CP (Figure S3 A-D). This demonstrates that in addition to a requirement for neuronal generation, *Ddx3x* also promotes neuronal migration between E13.5-E17.5. Aberrant neuron migration can be explained by cell-autonomous defects within migrating neurons, and/or by a defective radial glial scaffold. We assessed the radial glial scaffold using GFP to sparsely label radial glial fibers and anti-Laminin to label the basement membrane. In *Ddx3x-*deficient brains, the radial glial scaffold was intact with apparently normal connections of endfeet to the pia (Figure 5P). Given that DDX3X is expressed in both neurons and progenitors, this suggests it may influence neuronal migration cell autonomously. Altogether, these results establish critical functions of DDX3X in embryonic cortical development, and specifically in promoting differentiation of RGCs and IPs into neurons, and facilitating radial neuronal migration (Figure S3E).

### Wnt signaling is not affected in the developing neocortex

DDX3X has been reported to influence canonical Wnt signaling (Cruciat et al., 2013; Snijders Blok et al., 2015), which has known roles in progenitors and migrating neurons during corticogenesis (Bocchi et al., 2017; Harrison-Uy and Pleasure, 2012). Therefore, we examined Wnt signaling in *Ddx3x* LoF brains by expressing a canonical Wnt reporter composed of six TCF/Lef response elements driving expression of histone 2B (H2B):GFP (Figure S4A) (Ferrer-Vaquer et al., 2010). The fraction of Wnt reporter-positive cells among all cells electroporated at E14.5 with either scrambled or *Ddx3x* siRNAs was quantified at E17.5. We observed no significant difference in the fraction of Wnt reporter-positive cells between either condition, suggesting that, in this paradigm, *Ddx3x* LoF phenotypes are not due to deficient Wnt signaling (Figure S4B, p=0.905). This further indicates DDX3X may function directly in RNA regulation during these critical phases of brain development.

### *DDX3X* mutations induce RNA-protein cytoplasmic aggregates and alter translation

Having established LoF requirements of *Ddx3x* in cortical development, we next turned our attention to investigate the pathology of *DDX3X* missense mutations. These mutations contribute to about half of our clinical cohort and were associated with more severe clinical outcomes than LoF. DDX3X is an RNA binding protein, and promotes formation of RNA-protein stress granules both in response to cellular stresses and upon overexpression of native protein (Lai et al., 2008). Likewise, expression of cancer-associated mutations results in constitutive formation of stress granules (Valentin-Vega et al., 2016). Thus, we reasoned that *DDX3X* missense mutations may induce stress granule formation in neural progenitors and neurons in the absence of stress. This cell biological assay also enabled us to rapidly assess if phenotypic severity correlates with the most severe clinical outcomes.

To test these hypotheses, we selected 2 missense mutations from our cohort, one which showed mild outcomes but the most common recurrence (R376C) and one with severe clinical outcomes (R326H). We also tested 3 previously reported ID-associated mutations (R475G, R534H, P568L) (Snijders Blok et al., 2015), and a medulloblastoma-associated mutation previously shown to induce stress granules (G325E) (Valentin-Vega et al., 2016). The mutations were located in several protein domains and had variable predicted functional impact based on Polyphen-2 scoring (Figure S5A and Table S1). GFP-tagged human WT-*DDX3X* or mutant *DDX3X* were expressed in Neuro2A cells at equivalent levels (Figure 6A and S5B) and cells were fixed and imaged 24 hours later. WT-*DDX3X* expression resulted in primarily diffuse cytoplasmic localization with GFP-positive aggregates evident in just 5% of cells (Figure 6B). In contrast, expression of severe missense mutations caused aggregate formation in approximately 20% of transfected cells (Figure 7B and S5B, WT vs. R326H and R475G p=0.001). As a positive control, medulloblastoma-associated G325E also produced aggregates (Figure S5B, p=0.001), consistent with previous reports in HeLa cells (Valentin-Vega et al., 2016). Notably, expression of R376C, a recurrent mutation with a mild clinical outcome, did not significantly increase DDX3X aggregate formation relative to control (p=0.757). These data indicate that *DDX3X* missense mutations impair DDX3X subcellular localization resulting in aggregates, and suggest that aberrant aggregate formation may correlate with the severity of neurodevelopmental defect.

**Figure 6.**
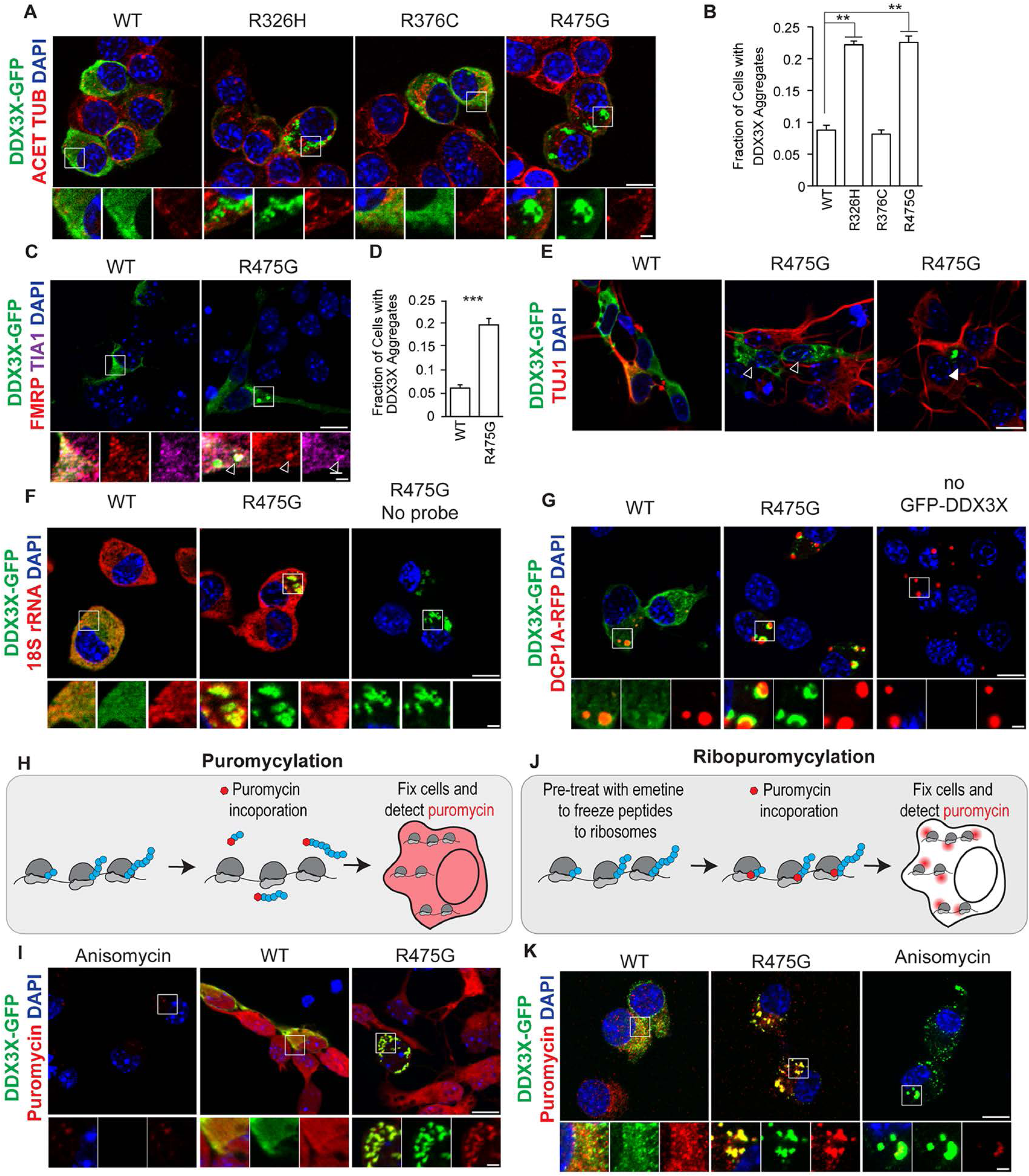
*DDX3X* missense mutations induce RNA-protein aggregates and impair translation. **A,** Images of Neuro2A cells transfected with DDX3X-GFP WT or point mutant constructs and stained for GFP (green), Acetylated-TUBULIN (red), and DAPI (blue) 24 hours after transfection. Higher magnification images from boxed regions are featured below. **B,** Quantification of percentage of transfected Neuro2A cells that contain DDX3X-GFP granules for WT and point mutant constructs counted 24 hours after transfection (n=3 biological replicates with at least 250 cells counted per replicate, One-way ANOVA p=1.057e-8, post-hoc Tukey test p value = 0.001 (WT vs. R326H, R475G); 0.757 (WT vs R376C)). **C,** Primary cortical cells transfected with WT or R475G DDX3X-GFP (green) and stained with FMRP (red) and TIA1 (magenta). R475G induces formation of granules that occasionally co-localize with FMRP and TIA1 (arrowhead). **D,** Quantification of percentage of transfected cells containing DDX3X-GFP granules in primary cortical cells 24 hours after transfection (n=3 biological replicates with at least 300 cells counted per replicate, Student’s t-test p=0.0006). **E**, Primary cortical cells transfected with WT or R475G DDX3X-GFP (green) and stained with TUJ1 (red). Both TUJ1-progenitors (empty arrowhead) and TUJ1+ neurons (filled arrowhead) contain DDX3X granules. **F,** Neuro2A cells transfected with DDX3X-GFP (green) and probed for 18S rRNA with smFISH probes (red). **G,** N2A cells transfected with DDX3X-GFP (green) constructs and DCP1A-RFP (red) to mark P-bodies. **H-I,** Puromycylation assay to monitor translation in primary cells 24 hours after transfection with WT or R475G-DDX3X (green). **H,** Schematic of puromycylation with ribosomes in gray, nascent peptide chains in blue, and puromycin in red. **I,** Puromycin signal (red) is blocked with the translation inhibitor anisomycin. **J-K,** Ribopuromycylation assay to visualize sites of active translation (red) in Neuro2A cells 24 hours after transfection of WT or R475G DDX3X-GFP (green). **J,** Schematic of ribopuromycylation with ribosomes in gray, nascent peptide chains in blue, and puromycin in red. **K**, Images of ribopuromycylation experiments in Neuro2A cells. Scale bars: 10 μm (upper panels), 2 μm (lower panels). Error bars = standard deviation.

We extended these studies to primary progenitors and neurons, and focused on R475G because of its robust aggregate formation phenotype in Neuro2A cells (Figure 6A and B). Equivalent levels of either WT or R475G DDX3X-GFP were expressed in primary cells derived from E14.5 dorsal cortices. Similar to Neuro2A cells, WT-DDX3X was primarily diffuse and cytoplasmic (Figures 6C and D). In contrast, R475G induced aggregate formation 3-fold more frequently with 19.8% of transfected cells containing aggregates compared to 6.1% for WT (p=0.0006). DDX3X aggregates were evident in both TUJ1-minus progenitor cells (empty arrowhead) and TUJ1-positive neurons (filled arrowhead) (Figure 6E). This indicates that a *DDX3X* missense mutation induces cytoplasmic aggregate formation in both neural progenitors and neurons.

We then used both Neuro2A cells and primary cortical cells to characterize the nature of these DDX3X aggregates. Cancer-associated *DDX3X* mutations induce formation of RNA-protein aggregates called stress granules (Valentin-Vega et al., 2016). Therefore, we investigated co-localization of DDX3X aggregates with canonical stress granule markers FMRP and TIA1 (Figure 6C). Notably, some, but not all, DDX3X aggregates co-localized with these canonical stress granule markers. We next determined if DDX3X aggregates contain RNA by coupling GFP staining with single molecule fluorescent *in situ* hybridization (smFISH) against 18s rRNA (Figure 6F). DDX3X aggregates co-localized with rRNA, suggesting these may be RNA-protein granules. Finally, we asked if DDX3X aggregates are processing bodies (P-bodies), which are cytoplasmic RNA-protein granules involved in mRNA decay and related to stress granules. Using the canonical P-body marker DCP1A (Decapping MRNA1A), we found DCP1A-RFP and DDX3X-GFP aggregates were often adjacent but did not co-localize (Figure 6F), suggesting that DDX3X aggregates are not P-bodies. Together, these data indicate that *DDX3X* missense mutations induce formation of cytoplasmic RNA-protein aggregates that share some features of stress granules.

To further characterize DDX3X aggregates, we examined the impact of expressing the R475G mutation upon global translation in E14.5 primary progenitors and neurons. Stress granules are typically associated with overall translational repression (Decker and Parker, 2012). Therefore, we predicted that if DDX3X aggregates are bona fide stress granules, overall translation should be reduced. We monitored protein synthesis using a puromycylation assay (Figure 6H), in which the tRNA analog puromycin is incorporated into nascent peptides and reflects global translation rates (Schmidt et al., 2009). Primary cortical cells transfected with equal amounts of WT or R475G DDX3X-GFP were pulsed with O-propragyl-puromycin for 20 minutes and then stained to detect puromycin incorporation (Figure 6I). Non-transfected cells and those expressing WT-DDX3X had diffuse and evenly distributed puromycin signal in the cytoplasm. This signal was blocked by anisomycin, a translational inhibitor, and by sodium arsenite, an oxidative stress that induces stress granules (Supplemental Figure 5C). Surprisingly, R475G cells showed focal accumulation of puromycin in aggregates. These data suggest that DDX3X RNA-protein aggregates contain newly synthesized proteins and are not translationally silent like stress granules.

The accumulation of newly synthesized proteins in DDX3X aggregates could be due to either local translation in granules or recruitment of nascent proteins following translation. To distinguish between these possibilities, we used ribopuromycylation (RPM) to visualize sites of translating ribosomes. This approach uses the translational inhibitor emetine to freeze puromycin-labeled peptides at the ribosome (Figure 6J) (David et al., 2012). The RPM signal was highly concentrated in DDX3X aggregates, and this signal was blocked by anisomycin (Figure 6K). These findings corroborate the puromycylation findings and demonstrate that translation can actively occur at DDX3X aggregates. While DDX3X may translationally repress some mRNAs, this contrasts with canonical characteristics of stress granules. Thus, based upon this translational signature and overlap with stress granule markers, we characterize DDX3X granules as RNA-protein aggregates with some stress-granule like features. Taken together, these results indicate that *DDX3X* missense mutations associated with neurodevelopmental pathology result in RNA-protein aggregates and impaired patterns of protein translation in neural progenitors and neurons.

### *DDX3X* mutations impair helicase activity that correlates with disease severity

Our cell biology experiments demonstrate that *DDX3X* mutations nucleate RNA-protein aggregates and disrupt DDX3X subcellular localization and translation patterns. This raises the question as to how *DDX3X* mutations induce these cell biological phenotypes. As *DDX3X* missense mutations fall almost entirely within the two helicase domains, we hypothesized they impair DDX3X helicase activity and RNA release following ATP hydrolysis. We further predicted that, similar to the observed formation of RNA-protein aggregates, if *DDX3X* mutations show impaired helicase activity, they might correlate with the degree of clinical severity. We therefore measured the ability of purified mutant DDX3 helicase-domain constructs to unwind RNA duplexes (Figure 7A) (Floor et al., 2016b). We selected recurrent *DDX3X* missense mutations associated with less severe clinical outcomes (R376C and I514T), severe impairment and RNA-protein aggregate formation (R475G), or with severe clinical impairment and PMG (T323I, R326H, I415del, T532M). Duplex unwinding by all mutant proteins was lower than WT by varying degrees (Figure 7B). Notably, there was mild to moderate slowing in the rate of unwinding with mutations found in less severely affected individuals (R376C and I514T), but complete loss of unwinding activity in all four of the PMG-associated mutations. The mild and severe phenotypes of R376C and R326H, respectively, correlate with cell biological phenotypes (as in Figure 6).

**Figure 7.**
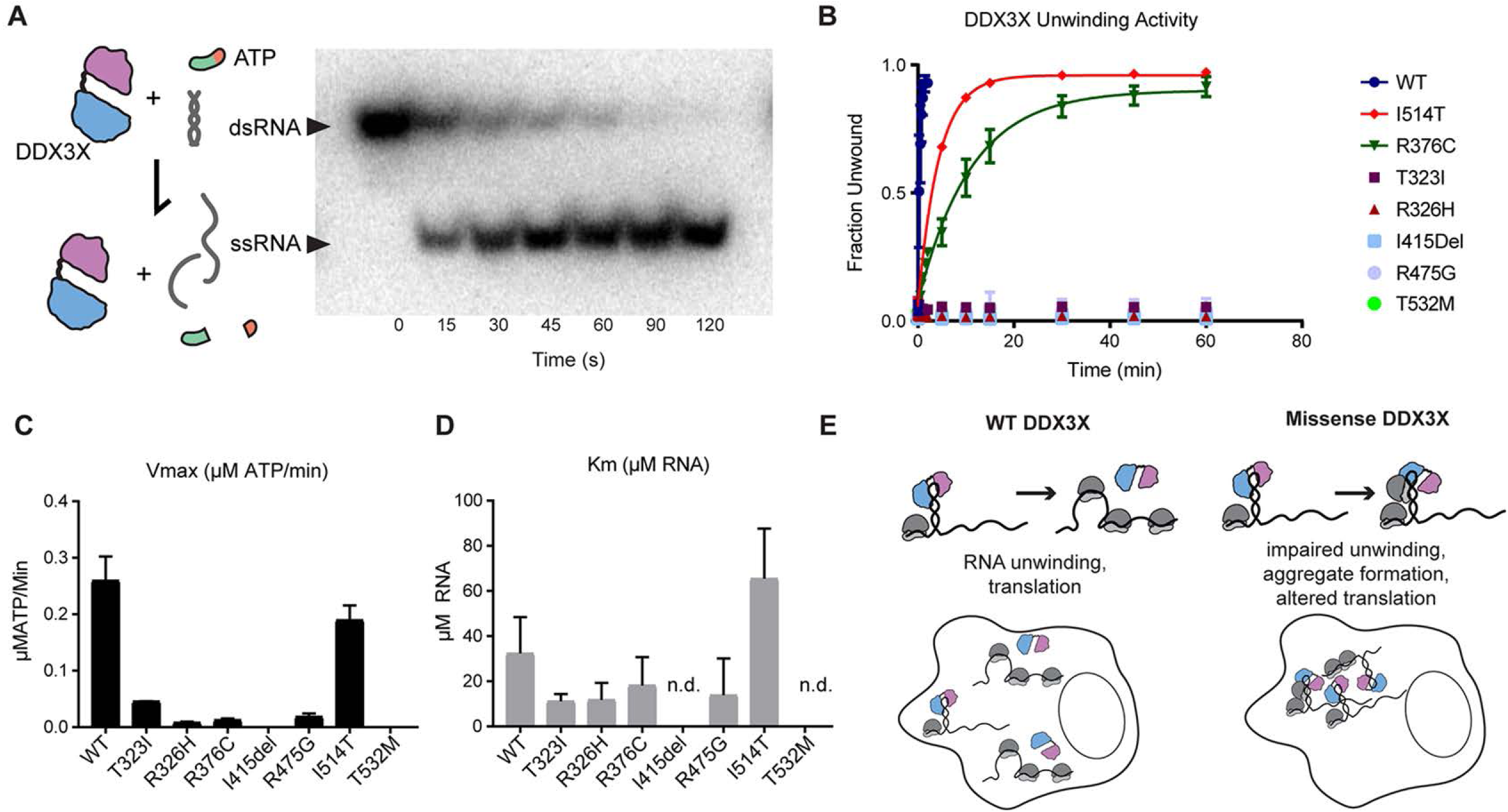
*DDX3X* missense mutants exhibit disrupted helicase activity. **A**, **Left,** Diagram of DDX3 activity tested in this assay. ATP is necessary for the initial binding of RNA, however ATP hydrolysis is not necessary for RNA unwinding but is required for release of the RNA from the protein complex. **Right**, blot of unwinding assay. DDX3 constructs were incubated with radioactively labeled RNA duplex and ATP, and aliquots at indicated time points were run on a non-denaturing gel (right). dsRNA runs more slowly than ssRNA, allowing percentage unwound to be quantified by band intensity**. B,** Unwinding assay for tested mutations in the first and second helicase domains WT, T323I, R326H, R376C, I415del, R475G, I514T, and T532, All mutants showed a decreased unwinding activity when compared to wild type (Far left). PMG mutants showed no unwinding activity while mutants found in patients with less severe clinical phenotypes (R376C, I514T) had less impaired unwinding. **C,** All mutants showed reduced Vmax when compared to the wild type. Two mutants (I415Del, T532M), both PMG mutants, had no detectable activity. **D,** Km for DDX3X mutants. I415del and T532M had unwinding curves which did not vary within the range of tested RNA concentrations (0-40 uM) so that Km could not be reasonably determined (n.d.). Majority of mutants had lower Km (indicating higher affinity for RNA) than the wild type with the exception of I514T. **E,** Proposed model based upon *in vitro* biochemical and cellular studies in which missense mutations in *DDX3X* found in our cohort impair unwinding and/or release from the RNA duplex, and induce aggregate formation, and ultimately alter translation in cells.

This impaired helicase activity may be the result of the DEAD box enzyme’s inability to bind RNA, to hydrolyze ATP and release the bound RNA, or both. To distinguish these possibilities, we further characterized the biochemical ATP hydrolysis activity and binding affinity of DDX3X to RNA by determining the V_max_ and K_m_ for DDX3X with respect to RNA concentration in the same group of mutants. We performed an ATP hydrolysis assay, in which both WT and mutant DDX3X peptide was incubated in a gradient of RNA concentrations with an excess of radioactively labelled ATP. ATP hydrolysis was measured through thin layer chromatography. Overall V_max_ and K_m_ were diminished in DDX3X mutants (Figures 7C and D). This suggests that specific missense mutations, especially those associated with PMG, exhibit a dominant negative effect of tighter binding to RNA (low K_m_) with reduced hydrolysis (low V_max_). The only exception to this trend was I514T, which had relatively poor unwinding activity, but also a considerably increased K_m_. This suggests a model consistent with haploinsufficiency in this patient with primarily significantly reduced binding to RNA, but also a mildly reduced unwinding activity. Taken together, these assays suggest that missense mutations can result in either dominant interfering interactions or a lack of functional interactions between DDX3X and RNA. These outcomes can result in a range of biochemical activity loss, which correlates with clinical and imaging-based disease severity.

### Discussion

*DDX3X* has recently been recognized as a frequently mutated gene in both ID and cancer, yet the mechanism by which *DDX3X* mutations impair brain development have not been defined. Here, we report the largest cohort of patients with *de novo* mutations in *DDX3X* (n=78) to date, highlighting corpus callosum abnormalities and PMG as two major brain anatomic phenotypes characteristic of *DDX3X* syndrome, with PMG strongly linked to missense mutations and to significant clinical impairment. Further, using *in vivo* mouse studies, we link these phenotypes to developmental mechanisms, showing that *DDX3X* is essential for proper cortical development by controlling neural progenitor differentiation and neuronal migration. Finally, using biochemical and cell biological assays, we show that *DDX3X* missense mutations disrupt helicase activity, resulting in RNA-protein aggregates and impaired patterns of translation. The severity of these biochemical and cellular impairments correlates closely with clinical severity, supporting the hypothesis that a subset of missense mutations function in a dominant negative manner, a novel and important finding for understanding DDX3X’s role in brain development. Altogether, our study uncovers new cellular and molecular mechanisms by which *DDX3X* mutations induce severe cortical malformations in humans, and broadly implicates new pathogenic etiologies of neurodevelopmental disorders.

### Location and type of mutation predict *DDX3X* clinical phenotype

In this study, we identify and characterize phenotypes for 78 patients, significantly extending our understanding of clinical outcomes associated with *DDX3X* mutations. Detailed analysis of our expanded cohort reveals the same recurrent *de novo* mutations in unrelated individuals result in markedly similar clinical phenotypes. Three pairs of patients (6 total) with PMG have mutations at recurrent amino acids, R326, I415, and T532. Indeed, the R326 mutation was also reported previously together with PMG (Snijders Blok et al., 2015). Within our cohort, we recognized 7 new PMG mutations (T198P, V206M, A233V, T323I, I415F, I415del, T532M). All of the patients with mutations at these amino acids had a more severe cerebral anatomic phenotype, including complete or partial absence of the corpus callosum. The correlation with PMG is important as these patients also tended to have more complicated clinical presentations, including epilepsy, autism spectrum disorder, severe intellectual disability, and structural cardiovascular malformations (ASD, PFO, PDA), than patients without PMG. Of the ten patients with PMG, nine were found to have missense mutations and one had a three-nucleotide deletion, which is predicted to result in a single amino acid deletion. This striking association between PMG and missense mutations supports the hypothesis that certain missense mutations have the potential to generate a more severe “dominant negative” phenotype. In contrast to the association of some missense mutations with more severe disease phenotypes, the most common recurrent mutation, R376C, showed a milder phenotype overall, with only 1 out of 4 patients being nonverbal past the age of 8, and all were normocephalic.

In contrast, none of the 32 patients with LoF mutations had PMG. Instead these patients with LoF mutations exhibited a milder spectrum of clinical phenotypes, including preservation of the ability to walk and talk. This degree of genotype-phenotype association was not evident in the first reported cohort of *DDX3X* patients (Snijders Blok et al., 2015), perhaps due to a smaller sample size.

### *Ddx3x* loss-of-function in mice impairs neuronal generation and migration

The malformations associated with *DDX3X* LoF, including microcephaly, abnormalities of the corpus callosum, and heterotopia, are predicted to result from perturbations during various stages of embryonic cortical development. Indeed, we demonstrate for the first time that *Ddx3x* LoF impairs essential processes of embryonic cortical development. First, we show that DDX3X is highly expressed in radial glial progenitors at key stages of neuron production and is essential for progenitor proliferation. As a result, acute knockdown of *Ddx3x* leads to an overall reduction in neuron number. How might *Ddx3x* impair neuron generation? One possibility is that DDX3X controls whether radial glia undergo proliferative versus neurogenic divisions. Another possibility, which is not mutually exclusive, is that DDX3X promotes cell cycle progression of RGCs and/or IPs and its depletion leads to more cycling progenitors. Indeed, DDX3X is implicated in cell cycle progression in mice and *Drosophila* (Chen et al., 2016; Li et al., 2014; Pek and Kai, 2011). How might this equate to clinical outcomes and anatomical malformations? Microcephaly, seen in patients carrying LoF mutations, is predicted to result from a generalized reduction of neurons. Moreover, the stages examined in this study are predicted to generate layer II/III neurons. Reduced superficial neurons may contribute to defects in corpus callosum development, seen in almost all of the patients (Fame et al., 2011).

Independent of its requirements in progenitors, DDX3X also promotes neuronal migration *in vivo*. As the radial glial scaffold was apparently intact following depletion of *Ddx3x* at mid-corticogenesis, we hypothesize that DDX3X promotes migration at least in part by acting cell-autonomously within neurons. Consistent with this interpretation, *Ddx3x* is strongly expressed in neurons, and previous studies demonstrate it is required for neurite outgrowth (Chen et al., 2016). We cannot rule out roles for DDX3X in RGC integrity prior to E13.5, however the electroporation paradigm should expose RGC defects. For example, injection of Cre into a *Igtb1*^lox/lox^ brain at E13.5 causes detachment of the radial glial fiber from the basement membrane by E14.5, arguing that acute defects can be induced at this stage (Radakovits et al., 2009). Regardless, future studies that utilize conditional *in vivo* mouse models to deplete *Ddx3x* specifically from neurons will be invaluable for understanding how DDX3X controls migration and for determining the extent to which migration defects are sustained postnatally. These phenotypes are particularly relevant as *DDX3X* mutations are associated with lamination alterations predicted to arise from aberrant migration. Moreover, neuronal requirements of DDX3X likely underlie clinical phenotypes of seizures and intellectual disability. Altogether, these mouse studies establish critical requirements of DDX3X in neuronal specification and migration, giving important clues as to how DDX3X LoF impairs cortical development.

### *DDX3X* missense mutations severely disrupt RNA metabolism

Our data pinpoint defective RNA metabolism as an underlying mechanism by which *DDX3X* missense mutations impact progenitors and neurons (Figure 7E). We show that *DDX3X* missense mutants have reduced helicase activity, predicted to impair release of the protein from RNA. Further we show that *DDX3X* missense mutants induce RNA-protein aggregates. Strikingly, *DDX3X* mutations associated with more severe clinical outcomes also induce stronger cell biological and biochemical phenotypes. As DDX3X normally forms functional trimers (Valentin-Vega et al., 2016), we predict that missense mutants which constitutively associate with RNA may function in a dominant negative fashion. Altogether this provides a molecular explanation for why more severe clinical phenotypes are associated with missense mutations than for carriers of nonsense and frameshift (presumed LoF) mutations.

But how might molecular defects induced by *DDX3X* mutations ultimately impact progenitors and neurons? In the case of missense mutations, some clues can be taken from studies of the RNA binding protein FUS, mutations in which cause the neurodegenerative disorders ALS and FTLD (Nolan et al., 2016). Similar to *FUS*, *DDX3X* missense mutations also induce RNA-protein aggregates, which only partially overlap with stress granule markers (Yasuda et al., 2017; Yasuda et al., 2013). Moreover, like FUS aggregates and in contrast to canonical stress granules, translation can occur at DDX3X aggregates. While it remains unclear whether protein products translated at these foci are functional, FUS is proposed to promote translation of some RNAs within aggregates, while repressing others, and we speculate the same occurs with DDX3X. In particular, *DDX3X* missense mutations could disrupt translation of key pro-proliferative or pro-migratory transcripts. These questions should be addressed in the future with comprehensive genomic studies of both LoF and missense variants.

Our study is amongst the first to link aberrant RNA-protein granules to the pathogenesis of neurodevelopment, which to date have been primarily associated with the etiology of neurodegeneration. One notable exception is the finding that *de novo* mutations in the RNA helicase *DHX30* induce stress granules and cause ID and developmental delay (Lessel et al., 2018). Indeed, RNA binding proteins, including RNA helicases, are highly expressed in the developing neocortex and many are essential for cortical development and associated with disrupted neurodevelopment (Lennox et al., 2018). Going forward, we argue that it will be important to consider widespread roles for aberrant RNA-protein aggregation in cortical development and disease. Our study therefore proposes a new paradigm for neurodevelopmental disorders.

In sum, by thoroughly exploring the biology behind phenotype-genotype correlates across a spectrum of patients, we have challenged assumptions about *DDX3X* disease etiology. Our study reinforces the need for using mutation-based approaches to understand the role of DDX3X in neurodevelopment. This potential to ultimately predict disease severity and clinical outcomes (both mild and severe) from biochemical or cell biological “read-outs” will be a benefit to patients and their parents both in terms of planning for the patient’s needs and for overall family planning.

## Materials and Methods

### Data Reporting

No statistical methods were used to predetermine sample size for this study. Recruitment for this study was not blinded.

### Sample

This study includes data from 78 participants from a network of seven clinical sites. Data collection sites had study protocols approved by their Institutional Review Boards (IRB), and all enrolled subjects had informed consent provided by parent/guardian.

### Data Collection Methods

The majority of mutations were identified through clinical whole exome sequencing either at referring and participating clinical centers or through the clinical genetic laboratories: GeneDx, Baylor, or BGI-Xome. Three patients (2839-0, 2839-3 and 2897-0) were ascertained as part of a larger study of genetic causes for Dandy-Walker malformation. Exome sequencing of 2839-0 and 2897-0 and their parents was performed as described (27264673, 28625504); 2839-3 is the identical twin of 2839-0 and Sanger sequencing confirmed she shared the mutation identified by exome sequencing in her sister. Genetic mutation data were obtained for all 78 participants. All clinical features and findings were defined by the medical records obtained from participants or from the completion of standardized behavioral and development measures. Thus, the parents of the probands were issued the following neuropsychological questionnaires according to their child’s age: Vineland Adaptive Behavior Scales, Second Edition (Vineland-II), Child Behavior Checklist (CBCL), Social Communication Questionnaire (SCQ) and Social Responsiveness Scale, Second Edition (SRS-2). 44 of the study’s 78 participants completed neuropsychological questionnaires. Comparison of population means for behavioral scales were performed using a Mann Whitney U-Test.

### MRI Review

This study includes MRI scans from 64 participants. All scans were initially reviewed locally by a radiologist for findings that, if present, were communicated to the participant and noted in the patient chart. DiCOM files of high quality MRI scans were then transmitted to the BDRP and neuroradiologic findings were noted in a standardized assessment of developmental features as previously utilized by our group (Hetts et al., 2006). All MRI findings were reviewed by a board certified pediatric neuroradiologist at UCSF blinded to genetic status as part of a larger more general review of potential agenesis of the corpus callosum cases.

### Plasmids and subcloning

The pX330-U6-Chimeric_BB-CBh-hSpCas9 was a gift from Feng Zhang (AddGene plasmid # 42230) (Cong et al., 2013). Guide sequences were designed using an online program (http://crispr.mit.edu/) and cloned into the pX330 vector as described by the Zhang lab (http://www.addgene.org/crispr/zhang/). The following guide RNA was used: *Ddx3x* exon 1 5’-AGTGGAAAATGCGCTCGGGC-3’. The TCF/LEF-H2B:GFP plasmid was a gift from Anna-Katerina Hadjantonakis (AddGene plasmid # 32610 (Ferrer-Vaquer et al., 2010)). Dcp1a-RFP was a gift from Stacy Horner’s lab. pCAG-DDX3X was generated by amplifying full length human *DDX3X* from cDNA and subcloning into the pCAG-EX2 vector using NEBuilder HiFi DNA Assembly Kit. GFP or FLAG tags were added upstream of the *DDX3X* to generate fusion proteins. The NEBuilder kit was also used to engineer point mutations.

All biochemical experiments were performed using DDX3X from amino acid residues 132-607 (NP-001357.3) in a pHMGWA vector backbone containing a 5’ 6xHis-MBP tag, as described, (Floor et al., 2016b). Mutant clones were generated using Quikchange XL-II site directed mutagenesis kit (Agilent Cat Number: 200521) with the following primers:

R376C 5’-CACATCGTATGGCAAACGCCTTTCGGTGGCAT-3’ R326H 5’-CATATCCACCAGGTGACCCGGGGTAGC-3’ T323I 5’-CGACCCGGGATAGCAACCAGCAAGTGG-3’ A233V 5’-ATCGGCAACAGAAAGACAGCCGTCTTACCGC-3’ T532M 5’-GCCGACACGACCCATACGACCAATACGGTGCACA-3’ I514T 5’-CCAATGTCAAACACGTGACCAACTTTGATTTGCCGAG-3’ A497V 5’-CGCCACTGCCGTCACCACCAGAATCGG-3’ R475G 5’-CACGCTGGCTGCCATCACCGTGAATGC 3’ I415M 5’-ACCTTCTGCGTCATATTCTCGCTAGTGGAGCCA-3’ I415del 5’-GTTGGCTCCACTAGCGAGAATACGCAGAAGGT-3’

### Purification of recombinant DDX3X WT and mutant

Plasmids encoding amino acid residues #132-607 were transformed into competent *E.coli* BL21-star cells, lysed by sonication in lysis buffer (0.5 M NaCl, 0.5% NP40, 10 mM Imidazole, 20 mM HEPES pH 7.5) and bound to nickel beads, as described (Floor et al., 2016b). Beads were washed with low salt (0.5M NaCl 20 mM Imidazole, 20 mM HEPES) and high salt wash buffers (1 M NaCl, 20 mM Imidazole, 20 mM HEPES pH 7.5) and eluted (0.5 M NaCl, 0.25 M imidazole, 100 mM Na_2_SO_4_, 9.25 mM NaH_2_PO_4_, 40.5 mM Na_2_HPO_4_). The 6xHis-MBP tag was cleaved overnight by *tev* protease at a 1:40 tev:protein ratio (w/w), and then dialyzed into ion exchange low salt buffer (200 mM NaCl, 20 mM HEPES ph7, 10% Glycerol, 0.5 mM TCEP). The sample was loaded onto a GE HiTrap heparin column (GE, 17-0406-01), eluted at 25% ion exchange high salt buffer (1 M NaCl, 20 mM HEPES pH 7, 10% Glycerol, 0.5 mM TCEP) and applied to Superdex 75 column (GE 17-0771-01) equilibrated in 500 mM NaCl, 10% Glycerol, 20 mM HEPES pH 7.5 and 0.5 mM TCEP. Peak fractions were collected, concentrated to ~50 μM, and then supplemented with 20% glycerol, diluted to 30 μM, and flash frozen in liquid nitrogen. Sample of the collected protein was run on a SDS-PAGE gel and stained with Coomassie to confirm protein purity and size.

### Radiolabelling and formation of RNA Duplex

Two complementary RNA molecules were synthesized by IDT: “5’ duplex” (5’-AGCACCGUAAGAGC-3’) and “3’ overhang (5’GCGUCUUUACGGUGCUUAAAACAAAACAAAACAAAACAAAA-3’). 5’ duplex RNA was labeled with ^32^P using T4 PNK (NEB, M0201S) for one hour at which point stop buffer was added to terminate the reaction. Labeled RNA was run on a 15% DNA denaturing gel, imaged on a phosphorscreen, and then bands were excised and RNA eluted overnight in 600 μL elution buffer (300 mM NaOAc pH 5.2, 1 mM EDTA, 0.5% SDS). Eluted RNA was then precipitated using 750 μL of 100% ethanol and immersed in dry ice for 1 hour 30 minutes and then resuspended in 20 μL DEPC H_2_O to > 200k cpm. RNA duplex was created by annealing 3 μL of 100 μM 3’ overhang RNA to 2.5 μL 5’ duplex of duplex RNA, heated to 95**°**C for 1 minute and allowed to cool to 30°C before purifying on a 15% native gel run at 8 watts.

### Duplex Unwinding Assays

This protocol was modified from Jankowski et al. (Jankowsky and Putnam, 2010). Radiolabeled, purified duplex RNA was diluted to ~333 cpm, and protein was diluted to 10 μM concentration. Reaction was run in 30 μL in reaction buffer (40 mM Tris-HCl pH 8, 5mM MgCl_2_, 0.1% IGEPAL, 20 mM DTT) with 3 μL of diluted RNA and 1 μL of 10 μM protein. 3 μL of 20 mM ATP:MgCl_2_ was added to initiate the reaction. At each time point 3 μL of reaction mix is removed and quenched in 3 μL of stop buffer (50 mM EDTA, 1% SDS, 0.1% Bromophenol blue, 0.1% Xylene cyanol, 20% glycerol). Reaction time points are then run on a 15% native polyacrylamide gel, at 5 watts for 30 minutes at 4°C. Gel is dried and imaged on a phospho-screen and quantified using ImageQuant software.

### ATP-ase activity Assay

The ATPase activity assay was performed with protein constructs previously purified and in the enzyme reaction buffer used the duplex unwinding assay reaction buffer, as described in Floor and Barkovich (Floor et al., 2016a). 100 μL of reaction master mix was made containing 3 μL of 10 μM DDX3X protein, 10 μL of 10x reaction buffer (400 mM Tris-HCl pH 8, 50mM MgCl_2, 1%_ IGEPAL, 200 mM DTT) and 26.9 μL of DEPC H_2_O. 2 μL of single stranded RNA was added to each 7 μL to create a dilution series of 0 μM, 0.625 μM, 1.25 μM, 2.5 μM, 5 μM, 10 μM, 20 μM and 40 μM. The single-stranded RNA was transcribed and gel purified, with sequence GGAAUCUCGCUCAUGGUCUCUCUCUCUCUCUCUCUCUCUCU. Radioactive ATP-(^32^P) (Perkin Elmer) was diluted to 666 μCi/uL. 1 μL of ATP at varying concentrations was added to initiate each reaction. At each time point 1 μL were spotted and quenched on PEI cellulose thin layer chromatography plates. Chromotography plates were run in a buffer of 0.5 M LiCl and 0.5 M Formic acid. Plates were then exposed to a GE Phosphoscreen for 1.5 hours and then imaged on a GE Typhoon imager.

### Mouse husbandry

All animal use was approved by either the Division of Laboratory Animal Resources from Duke University School of Medicine or by the University of Queensland Animal Ethics committee in accordance with the Australian Code of Practice for the Care and Use of Animals for Scientific Purposes. C57BL/6J mouse embryos of either sex (Jackson Laboratory) were used for all experiments, except for coronal section *in situ* hybridizations (Figure 3 F-I) and postnatal immunofluorescence (Figure S2), which both used CD1 mice. Plug dates were defined as E0.5 on the morning the plug was identified.

### siRNAs

Mouse *Ddx3x* siRNAs were obtained from Qiagen. siRNA experiments were performed by pooling 4 siRNAs with the following target sequences: 5’-CTGATAATAGTCTTTAAACAA-3’, 5’-TCCATAAATAATATAAGGAAA-3’, 5’-CTCAAAGTTAATGCAAGTAAA-3’, 5’-CACAGGTGTGATACAACTTAA-3’.

### Cell lines and primary cultures

Neuro2A cells were cultured in DMEM supplemented with 10% FBS and 1% penicillin/streptomycin. Neuro2A cells were transfected with Lipofectamine 2000 (Invitrogen). For the DDX3X granule formation assay, equal amounts of plasmid were transfected for each construct and cells were fixed 24 hours after transfection. Samples were blinded prior to quantification on the microscope. Primary cortical cultures were derived from E12.5 – E14.5 embryonic dorsal cortices, as previously described (Mao et al., 2015). Microdissected tissue was dissociated with 0.25% Trypsin-EDTA + DNase for 5 minute at 37°C before trituration with a pipette. Cells were plated on poly-D-lysine coated coverslips in 12 well culture plates. Cells were grown in DMEM supplemented with B27, N2, N-acetyl-L-cystine, and bFGF. Primary cortical cells were transfected with the Amaxa Nucleofector in P3 solution.

### Western blot and qRT-PCR anlaysis

Neuro2A cells were transfected with either Cas9 with or without *Ddx3x* sgRNA or scrambled or *Ddx3x* siRNAs. Protein was harvested 72 hours after transfections in RIPA lysis buffer with protease inhibitors. Lysates were run on 4-20% mini-PROTEAN TGX precast gels. Gels were transferred to PVDF membranes, blocked with 5% milk/TBST, probed primary antibodies overnight at 4°C, and secondary HRP-conjugated antibodies at room temperature for 1 hour. The following primary antibodies were used: mouse anti-DDX3X (Santa Cruz, sc-365768, 1:100) or mouse anti-Tubulin (Sigma, T6199, 1:1000). Blots were developed with ECL and quantified by densitometry in ImageJ. RNA was extracted from transfected Neuro2A Cells using TriReagent (Sigma), followed by cDNA synthesis with iScript Reverse Transcriptase (BioRad). The primers used for qRT-PCR are as follows: *Ddx3x* forward 5’-TGGAAATAGTCGCTGGTGTG-3’ and reverse 5’-GGAGGACAGTTGTTGCCTGT-3’; *Actb* forward 5’-AGATCAAGATCATTGTCCT 3’ and reverse 5’ CCTGCTTGCTGATCCACATC 3’.

### In utero electroporation

Plasmids were delivered to embryonic brains as previously described (Mao et al J Neuro 2015). Briefly, E13.5 or E14.5 embryos were injected with 1-1.5 uL of plasmid DNA mixed with Fast Green Dye. Plasmids were used at the following concentrations: pCAG-GFP (1.0 ug/uL), pX330 empty or pX330-*Ddx3x* Ex1 sgRNA (2.4 ug/uL), TCF/LEF-H2B:GFP (1.25 ug/uL), pCAG-mCherry (1.0 ug/uL). Scrambled or *Ddx3x* siRNAs were injected at 1 μM. Following injection, embryos were pulsed five times with 50 V for 50 ms. Embryonic brains were harvested 48-72 hours later.

### In situ Hybridization

*In situ* hybridization was performed as described in Moldrich et al, 2010. The *Ddx3x* riboprobe was generated in-house using the primers corresponding to those used in the Allen Developing Mouse Brain Atlas (Website: © 2015 Allen Institute for Brain Science. Allen Developing Mouse Brain Atlas [Internet]). Available from: http://developingmouse.brain-map.org): Forward primer: 5’ AAGGGAGCTCAAGGTCACAA 3’, Reverse primer: 5’ CCTGCTGCATAATTCTTCC 3’. Using mouse cortex cDNA, these primers were used to amplify a 908 base pair fragment. This fragment was purified and cloned into pGEM^®^-T Vector System (Promega). The plasmid was then linearized (SacII restriction enzyme, New England BioLabs), purified (PCR Clean up Kit, Qiagen), transcribed (Sp6 RNA Polymerase, New England BioLabs) and digoxigenin-labelled (DIG RNA labelling Mix, Roche) to generate the riboprobe. *In situ* hybridization against *Ddx3x* mRNA was performed on 20 µm cryostat sections for embryonic stages and 50 µm vibratome sections for postnatal brains in WTCD1 mice.

### Immunofluorescence

Embryonic brains were fixed overnight in 4% PFA at 4°C, submerged in 30% sucrose overnight, and embedded in NEG-50. 20 μm frozen sections were generated on a cryotome and stored at - 80. Sections were permeabilized with 0.25% TritonX-100, blocked with either 5% NGS/PBS or MOM block reagent (Vector Laboratories) for 1 hour at room temperature. Sections were incubated with primary antibodies overnight at 4°C, and secondary antibodies at room temperature for 30-60 minutes. Cultured cells were fixed for 15 minutes at room temperature with 4% PFA, permeabilized with 0.5% Triton X-100, blocked with 5% NGS, incubated with primary antibodies for 1 hour at room temperature and secondary antibodies for 15 minutes at room temperature. Images were captured using a Zeiss Axio Observer Z.1 equipped with an Apotome for optical sectioning. The following primary antibodies were used: mouse anti-DDX3 (Santa Cruz, sc-365768, 1:100), rabbit anti-DDX3X (Protein Tech, 11115-1-AP, 1:150), mouse anti-TUJ1 (Biolegend, 801202, 1:1000), mouse anti-NESTIN (BD Biosciences, BD401,1:100), rabbit anti-PAX6 (Millipore, AB2237, 1:1000), rabbit anti-TBR2 (Abcam, AB23345, 1:1000), rabbit anti-CC3 (Cell Signaling, 9661, 1:250), rabbit anti-NeuroD2 (Abcam, AB104430, 1:500), chicken anti-GFP (Abcam, Ab13970, 1:1000), rabbit anti-Laminin (Millipore, AB2034, 1:200), rabbit anti-acetylated Tubulin (Sigma, T7541, 1:500), rabbit anti-FMRP (Sigma, F1804,1:500), mouse anti-TIA1 (Abcam, AB2712, 1:100), mouse anti-puromycin (DSHB, PMY-2A4, 1:100), anti-RFP (Rockland, 600-401-379S, 1:500), guinea pig anti-NeuN (Merck, ABN90P, 1:1000), chicken anti-GFAP (Abcam, ab4674, 1:1000), goat anti-OLIG2 (Santa Cruz, sc19969, 1:200), goat anti-IBA1 (Abcam, ab5076, 1:1000), mouse anti-FOXJ1 (eBioscience, 149965-82, 1:1000), and rabbit anti-DDX3X (Sigma Aldrich, HPA001648, 1:500). Secondary antibodies used in embryonic brains and cell culture experiments were Alexa Fluor-conjugated (Thermo Fisher Scientific, 1:500). Those used in postnatal brains were Alexa Fluor-conjugated secondary antibodies (Thermo Fisher Scientific), biotinylated-conjugated secondary antibodies (Jackson Laboratories) used in conjunction with AlexaFluor 647-conjugated Streptavidin (Thermo Fisher Scientific) and CF dyes conjugated secondary antibodies (Biotium). Postnatal brains were collected and fixed as previously described (Moldrich et al., 2010 and Piper et al., 2011), then post-fixed with 4% paraformaldehyde (ProSciTech) for 2 to 4 days and stored in 1x Dulbecco’s phosphate buffered saline (Lonza) with 0. 1% sodium azide (Sigma Aldrich). Brains were sectioned on a vibratome (Leica) at 50 µm thickness in coronal orientation. Fluorescence immunohistochemistry was performed as in Chen et al., 2017.

### Quantification and Binning Analysis

All mouse embryo experiments were blinded throughout processing, imaging, and analysis. For the binning analysis, 450 μm wide cortical columns were broken down into 5 evenly spaced bins spanning from the ventricular to the pial surface. Each GFP+ cell was assigned to a bin to calculate the distribution. At least 3 sections were analyzed per embryo, with multiple embryos per condition (see figure legend for exact sample size).

### Puromycylation, Ribopuromycylation, and FISH

For puromycylation, cultured cells were pre-treated with 40 μM anisomycin or 0.5 mM sodium arsenite for 10 minutes, followed by a 20 minute incubation with 10 μM O-propargyl-puromycin (Life Technologies).. Cells were rinsed with PBS, and fixed in 4% PFA for 15 minutes, and puromycin was conjugated to AlexaFluo594 with Click-It Technology according to the manufacturer’s protocol. For ribopuromycylation, cultured cells were pre-treated with 208 μM emetine for 15 minutes, then pulsed with 91 μM puromycin for 5 minutes in the presence of emetine. Anisomycin used at 40 μM as a competitive inhibitor of puromycin. Cells were extracted with cold polysome buffer (50 mM Tris-HCl pH 7.5, 5 mM MgCl_2_, 25 mM KCl, 355 μM CHX, EDTA-free protease inhibitors, and 10 U/mL RNaseOut) containing 0.015% digitonin on ice for 2 minutes, rinsed with cold polysome buffer, then fixed with 4% PFA for 15 minutes at room temperature. Puromycin incorporation was detected using the 2A4 anti-PMY antibody (DSHB) at 1:100, and imaged with fluorescent microscopy. Stellaris single molecule FISH probes against 18S rRNA were purchased from BioSearch Technology. Cells were fixed with PFA for 10 minutes, permeabilized with 70% EtOH for 1-2 hours at 4°C, rinsed with wash buffer (10% formamide,2x SSC), hybridized with probe at 1:100 overnight at 37°Cin hybridization buffer (100 mg/mL dextran sulfate, 10% formamide, 2x SSC), and rinsed again with buffer before mounting.

## Acknowledgements

We thank the patients, their families, and referring physicians for their important contribution to our ongoing work on these disorders. We thank members of the Silver lab for helpful discussions and reading the manuscript. This work was supported by the Holland-Trice Foundation (to DLS); NIH R01NS083897 (to DLS); NIH F31NS0933762 (to ALL); NIH 1R01NS058721 (to EHS, WBD, and LJR); NIH 5R01NS050375 (to WBD); the DDX3X Foundation (to EHS), the Principal Research Fellowship from the National Health and Medical Research Council, Australia (to LJR); the UCSF Program for Breakthrough Biomedical Research, funded in part by the Sandler Foundation (to SNF); the California Tobacco-Related Disease Research Grants Program 27KT-0003 (to SNF); the Dandy-Walker Alliance (to KAA and WBD); the HUGODIMS consortium RC14_0107, funded in part by the French Ministry of Health and the Health Regional Agency from Poitou-Charentes (to SK); and Frédérique Allaire from the Health Regional Agency of Poitou-Charentes (to SK). The content is solely the responsibility of the authors and does not necessarily represent the official views of the funding sources.

## Author contributions

ALL, RJ, DLS and EHS conceived of and designed the study, analyzed all data, and wrote the manuscript. ALL and CM, with guidance from JB, carried out mouse expression analyses. ALL, with assistance from CJS, conducted all mouse functional studies, and all cell biological (RNA-protein aggregate and translation) assays of DDX3X function. RJ conducted the biochemical studies with guidance from SNF. AJB and EHS reviewed and scored all human brain imaging. LJR, WBD, and AJB contributed to clinical study design and analysis of human imaging. IL conducted clinical statistical analyses. The following authors participated in patient recruitment: LS, BF, KAA, AA, DBV, SB, PRB, LB, PC, BHYC, BC, SD, NDD, LF, DH, AMI, BI, BK, AK, EWK, PK, SK, DMC, CM, CN, DR, LNB, CT, JT, MV, AZ. All authors edited the manuscript.

---

**Supplemental Figure 1.**
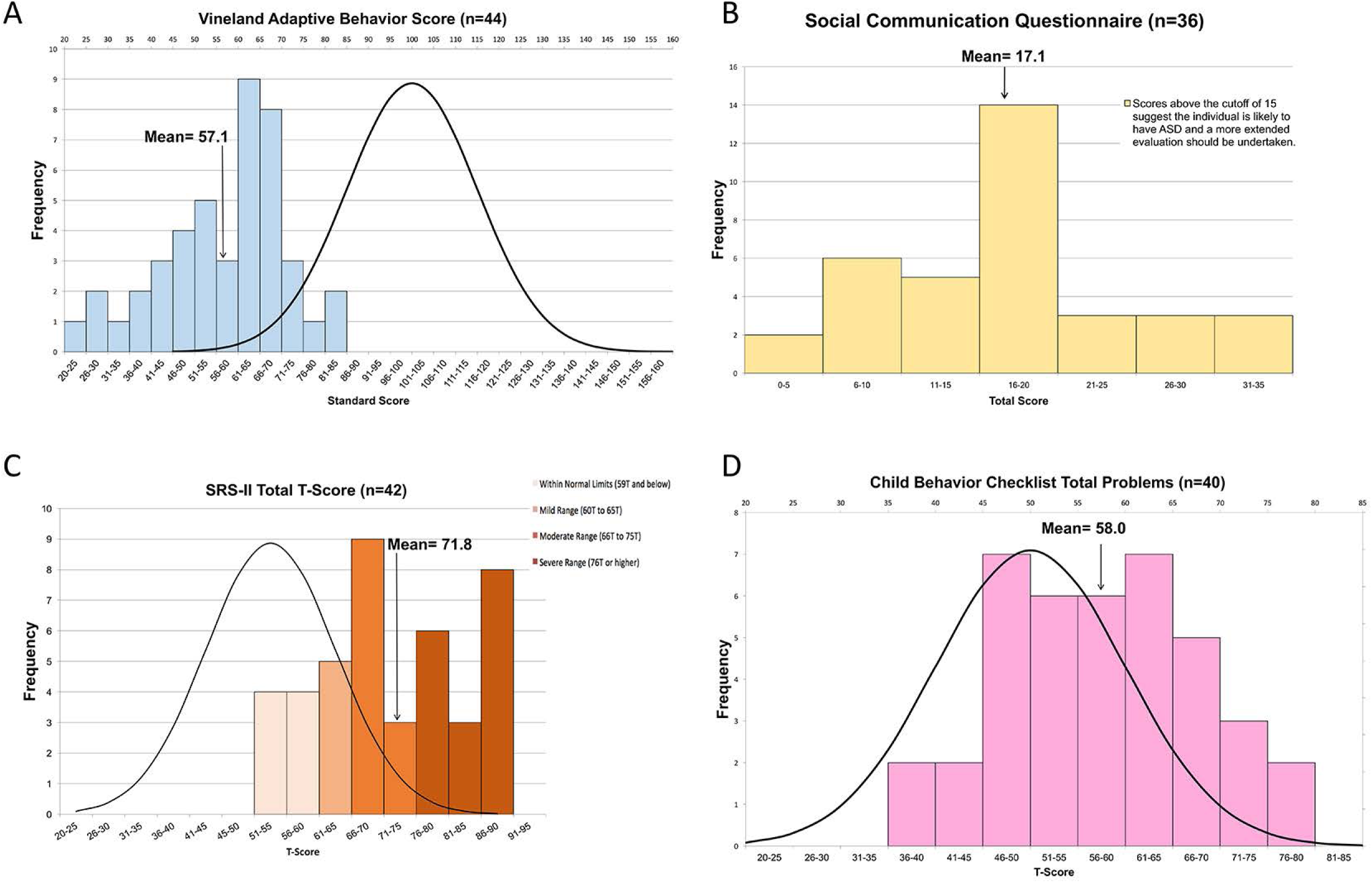
Scores on four measures of clinical assessment: **A.** the Vineland Adaptive Behavior Scales (VABS), **B.** the Social Communication Questionnaire (SCQ), **C.** the SRS-II, and **D.** the Child Behavioral Checklist (CBCL). If a standard curve for neurotypical individuals was available (VABS, SRS-II, and CBCL) it was displayed alongside the binned data for the DDX3X participants. The number of individuals for whom data was available for each test is indicated in the title and the graph also indicates the mean for the DDX3X cohort. There is a significant difference between DDX3X patients and neurotypical controls on all of these measures. For example, the VABS score had a mean of 57.1, which is nearly three standard deviations below the mean for controls (mean = 100).

**Supplemental Figure 2.**
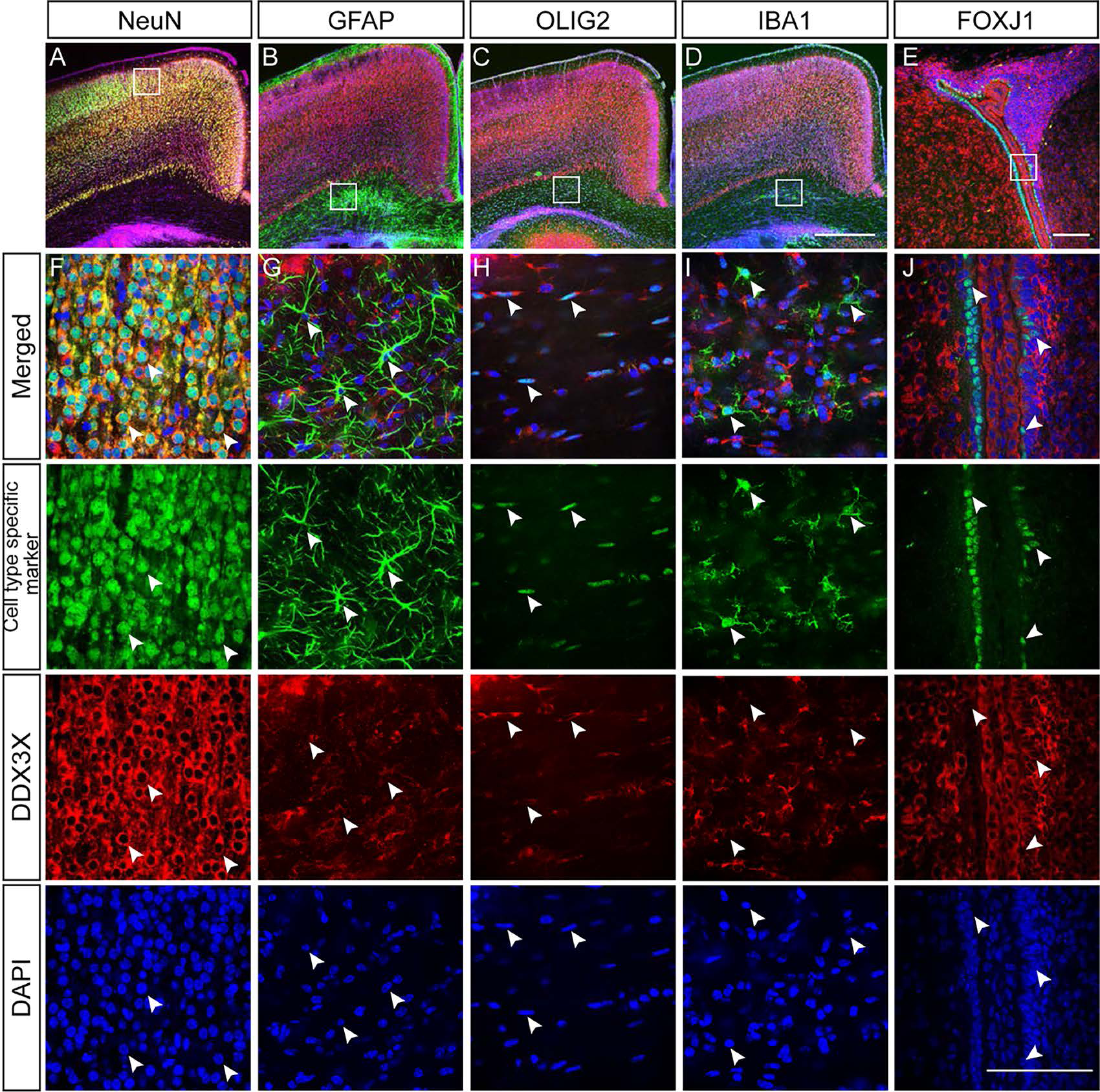
Immunohistochemistry depicting DDX3X expression in postnatal brains. **A-E,** Immunohistochemistry on coronal vibratome sections of P5 CD1 brains (n = 4) with antibodies against DDX3X (red) and indicated specific cell type markers NeuN (neurons), GFAP (astrocytes), OLIG2 (oligodendrocytes), IBA1 (microglia) or FOXJ1 (ependymal cells). **F-J,** Low power images (20x) showing parts of the neocortex and corpus callosum where cingulate cortex is to the right. (I) Low power image (20x) showing ependymal cells lining the lateral ventricle and choroid plexus within the ventricle. High power images (60x) for the insets (boxed regions) respectively. Arrow heads represent examples of co-expression. Scale bar = 500 µm for (A-E); 100 µm for (F-J).

**Supplemental Figure 3.**
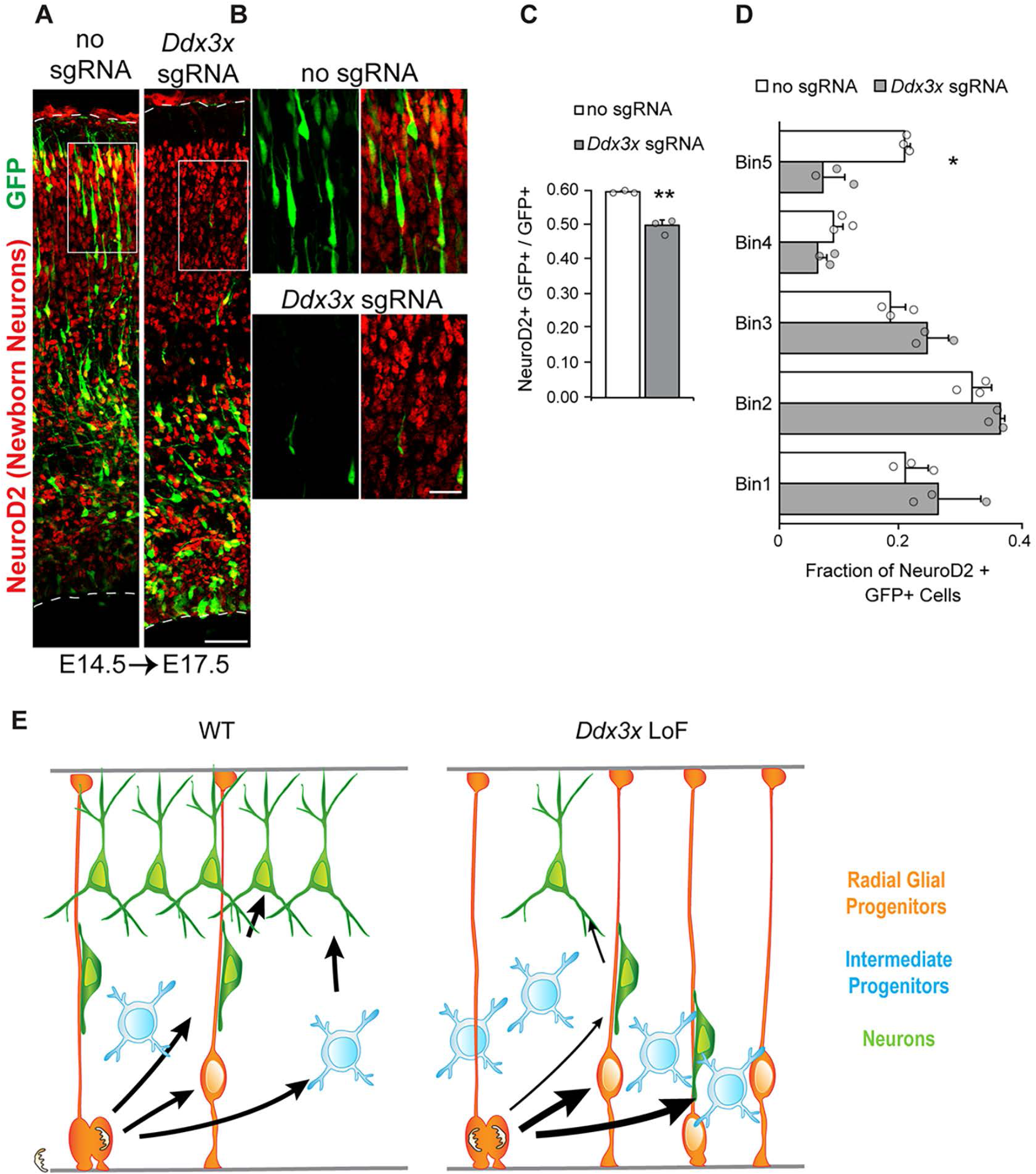
*Ddx3x* is required for neuron generation and neuron migration from E14.5 to E17.5. **A-B,** Sections of electroporated E14.5 – E17.5 brains stained with GFP (green) and NeuroD2 (red). Boxed regions in CP in **A** are shown at higher magnification in **B. C**, Quantification of percentage of GFP positive cells expressing NeuroD2 (n=3 embyos per condition, Student’s *t-*test p=0.0075). **D,** Quantification of the distribution of GFP+ NeuroD2+ cells in **A** (n=3 embryos per condition, Student’s *t*-test p=0.3252(Bin1); 0.1242(Bin2); 0.0783(Bin3); 0.1050(Bin4); 0.0179(Bin5)). **E,** Cartoon representation of phenotypes associated with *Ddx3x* LoF, inlcuding increased progenitors, fewer neurons and impaired neuronal migration. Scale bars: 50 μm (**A**); 15 μm (**B**). Error bars= SD.

**Supplemental Figure 4.**
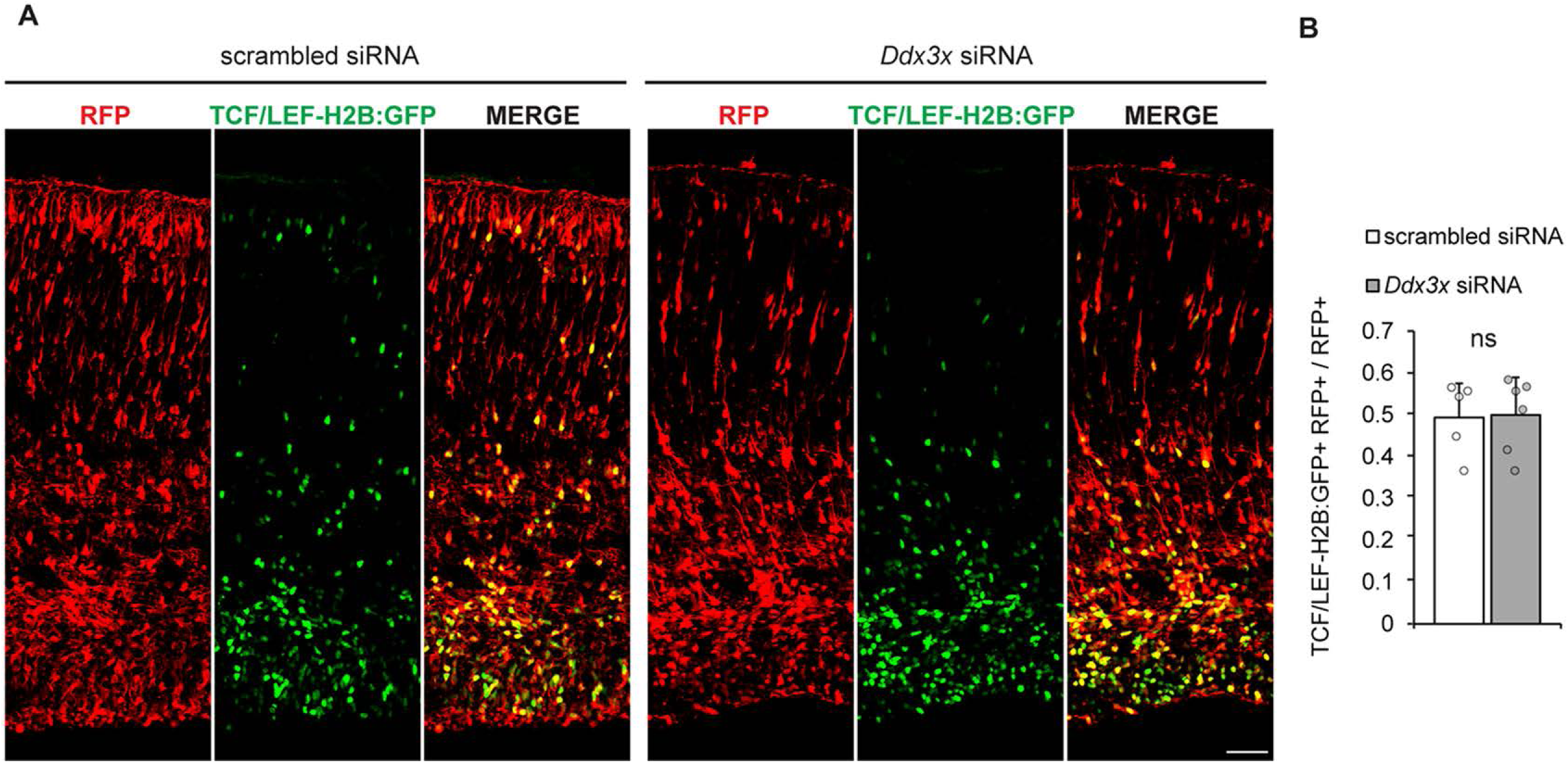
*Ddx3x* is dispensable for Wnt signaling in the developing brain. **A**, Representative images of coronal sections of E17.5 brains *in utero* electroporated at E14.5 with pCAG-mCherry, TCF/LEF-H2B:GFP, and scrambled or *Ddx3x* siRNAs. Sections were stained with anti-RFP (red) and anti-GFP (green). **B,** Quantification of percentage of RFP+ cells also positive for TCF/LEF H2B:GFP (n=5 embryos (scrambled) or 6 embryos (*Ddx3x*), Student’s *t*-test p=0.9045). Scale bars: 50 μm (**A**). Error bars = SD.

**Supplemental Figure 5.**
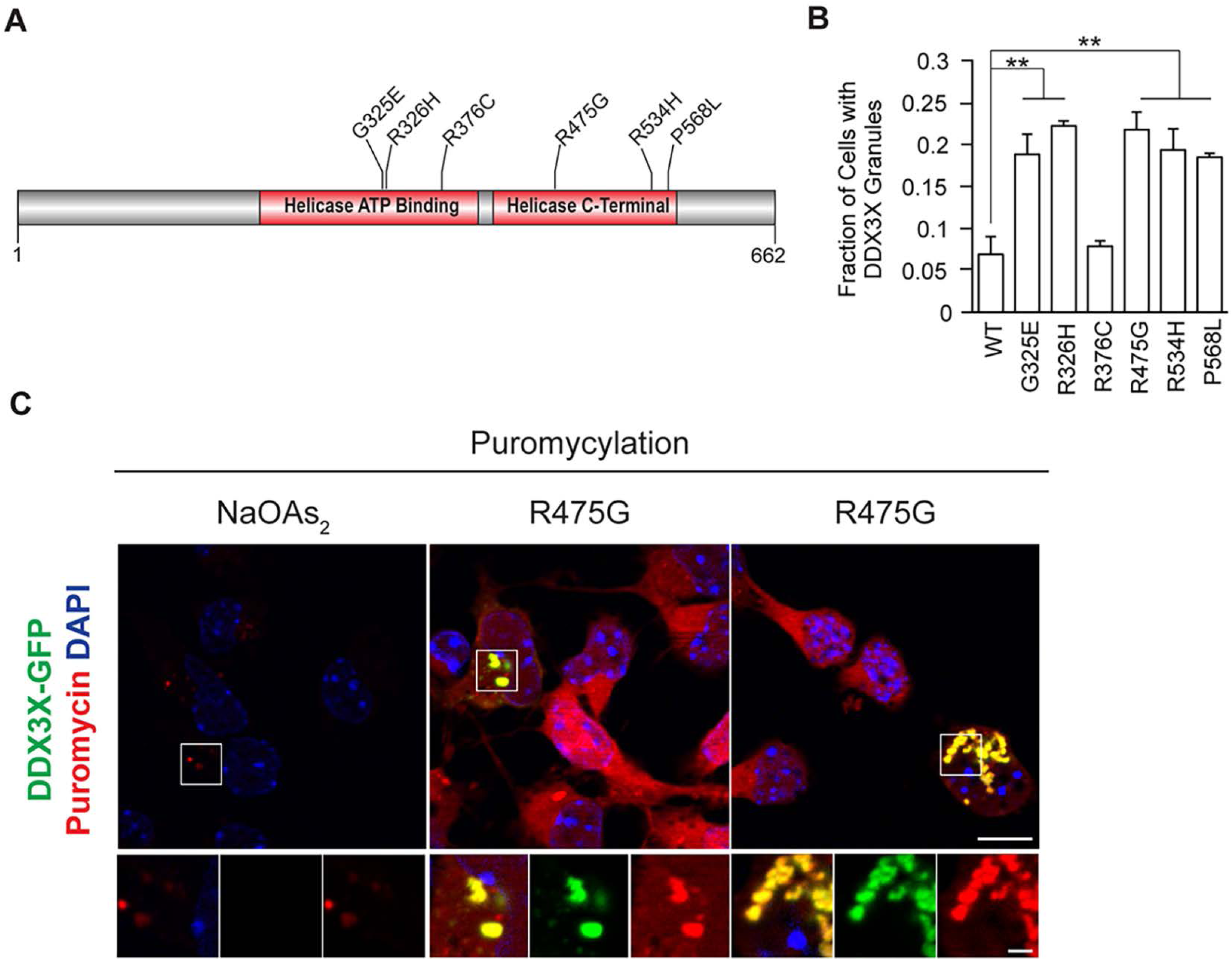
*Ddx3x* missense mutations induce protein aggregates and impair translation. **A**, Schematic of DDX3X structure with missense variants tested in the cell culture experiments. **B,** Expanded graph, related to Figure 7B, of quantification of percentage of transfected N2A cells that contain GFP-DDX3X granules for WT and point mutant constructs counted 24 hours after transfection (n=3-6 biological replicates with at least 250 cells counted per replicate, One-way ANOVA p=1.9051e-13, post-hoc Tukey test p value = 0.001 (WT vs. G325E, R326H, R475G, R534H, P568L); p value = 0.899 (WT vs R376C)). **C,** Puromycin incorporation assay to monitor translation in primary cells 24 hours after transfection with R475G GFP-DDX3X (green). Puromycin signal (red) is blocked with the stress granule-inducing agent sodium arsenite (NaOAs_2_).

